# Oxidative stress and lipidocentric mechanism of hyperglycemic glycolaldehyde damage to skeletal muscle cells

**DOI:** 10.64898/2026.11.23.767857

**Authors:** Savannah E. Banks, Aayush Vishwanath, Vijay Kumar, Thomas J. Hund, Narasimham Parinandi

## Abstract

Currently, diabetes-induced skeletal muscle complications affect over half of diabetic patients yet the pathway by which this hyperglycemic damage occurs has yet to be fully elucidated. As such, the goal of this study is to investigate the hypothesis: Glucose-derived glycolaldehydes, glyoxal and methylglyoxal, formed in hyperglycemia cause damage to the skeletal muscle myoblasts through oxidative stress and formation of pro-inflammatory eicosanoids upon activation of phospholipase A_2_ (PLA_2_) and cyclooxygenase (COX). To test this hypothesis skeletal muscle myoblasts, C2C12s, were exposed to various concentrations of glyoxal and methylglyoxal then tested for metabolic activity, reactive oxygen species production, redox status, ATP production, membrane integrity, cellular proliferation, enzyme activity, and loss of mitochondrial lipids. Doses of the agonist at 1 mM and 2 mM showed significant changes in all tested areas and were significantly attenuated by NAC, Ibuprofen, AACOCF3 and metformin. For the first time, these results suggest a plausible pathway by which diabetes-induced skeletal muscle deterioration occurs and identifies possible therapeutic endpoints.

## Introduction

According to the CDC, it is estimated that roughly 40.1 million Americans are afflicted with diabetes (diagnosed or undiagnosed) or around 12% of the national population^1^. This condition is primarily characterized by excess blood sugar, hyperglycemia, resulting from decreased efficiency or quantity of insulin produced in the body. This condition greatly affects morbidity rates and quality of life for patients afflicted, so clear understanding of the various pathways is crucial towards progress to effective therapeutics.

The accumulated glucose then undergoes multiple oxidation steps through glycolysis producing the glycolaldehyde Methylglyoxal (MGO) .^2^ An additional glycolaldehyde, Glyoxal (GO), can be produced through the Amadori rearrangement of a Schiff base and the subsequent autooxidation of the Amadori product. The structures of both carbonyls are shown in figure 1. ^3^

**Figure 1.**
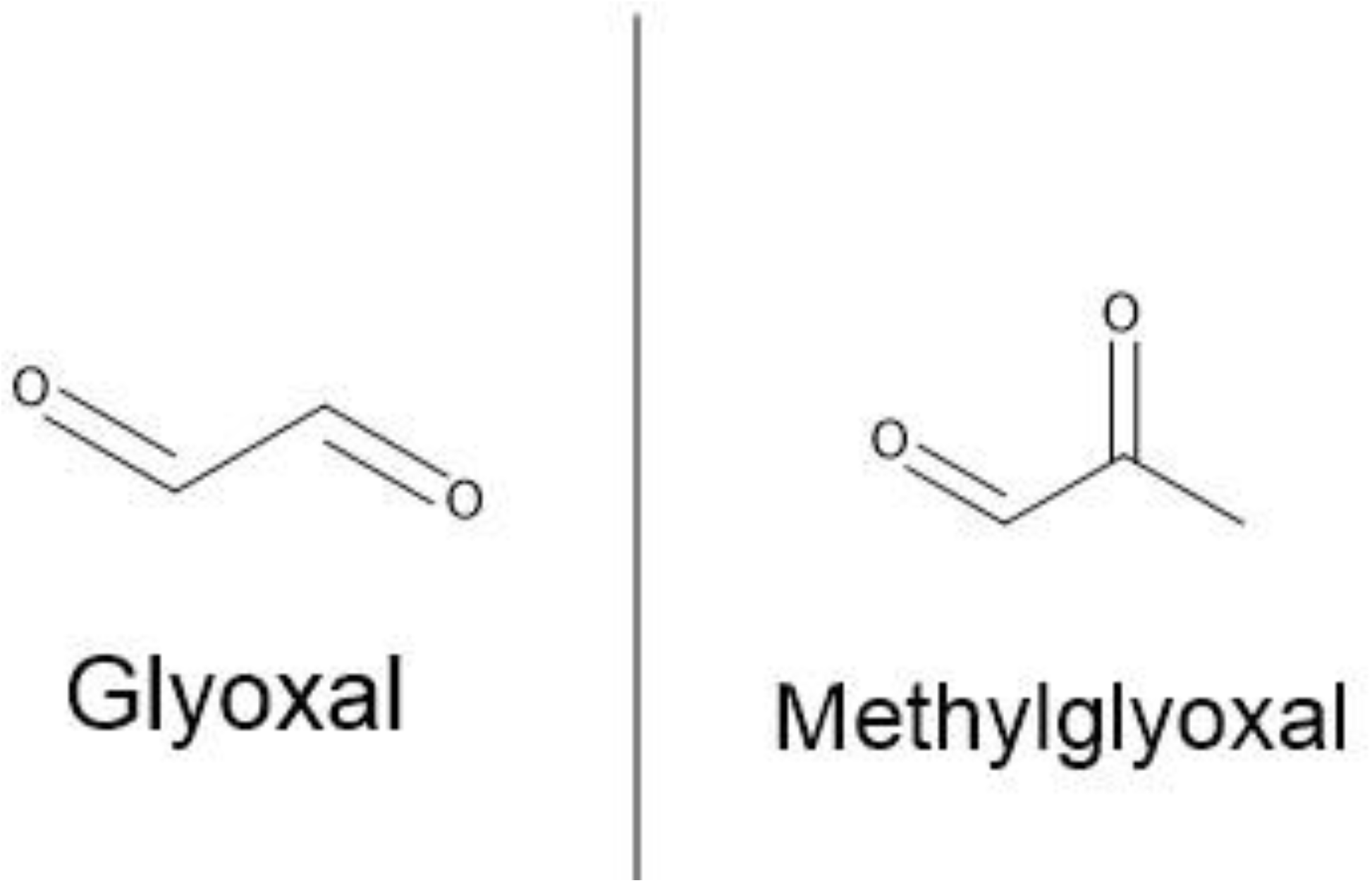
MGO and GO Structure Structural representation of the reactive dicarbonyls glyoxal (left) and methylglyoxal (right).

MGO and GO are key players in protein glycation and through interactions with proteins, nucleic acids, and peptides, MGO and GO form advanced glycation end products (AGEs). As shown in the simplified pathway in figure 2, common AGE produced by methylglyoxal is Nε-(1-carboxyethyl) lysine (CEL), whereas glyoxal produces Nε-(carboxymethyl)lysine (CML). ^4^

**Figure 2.**
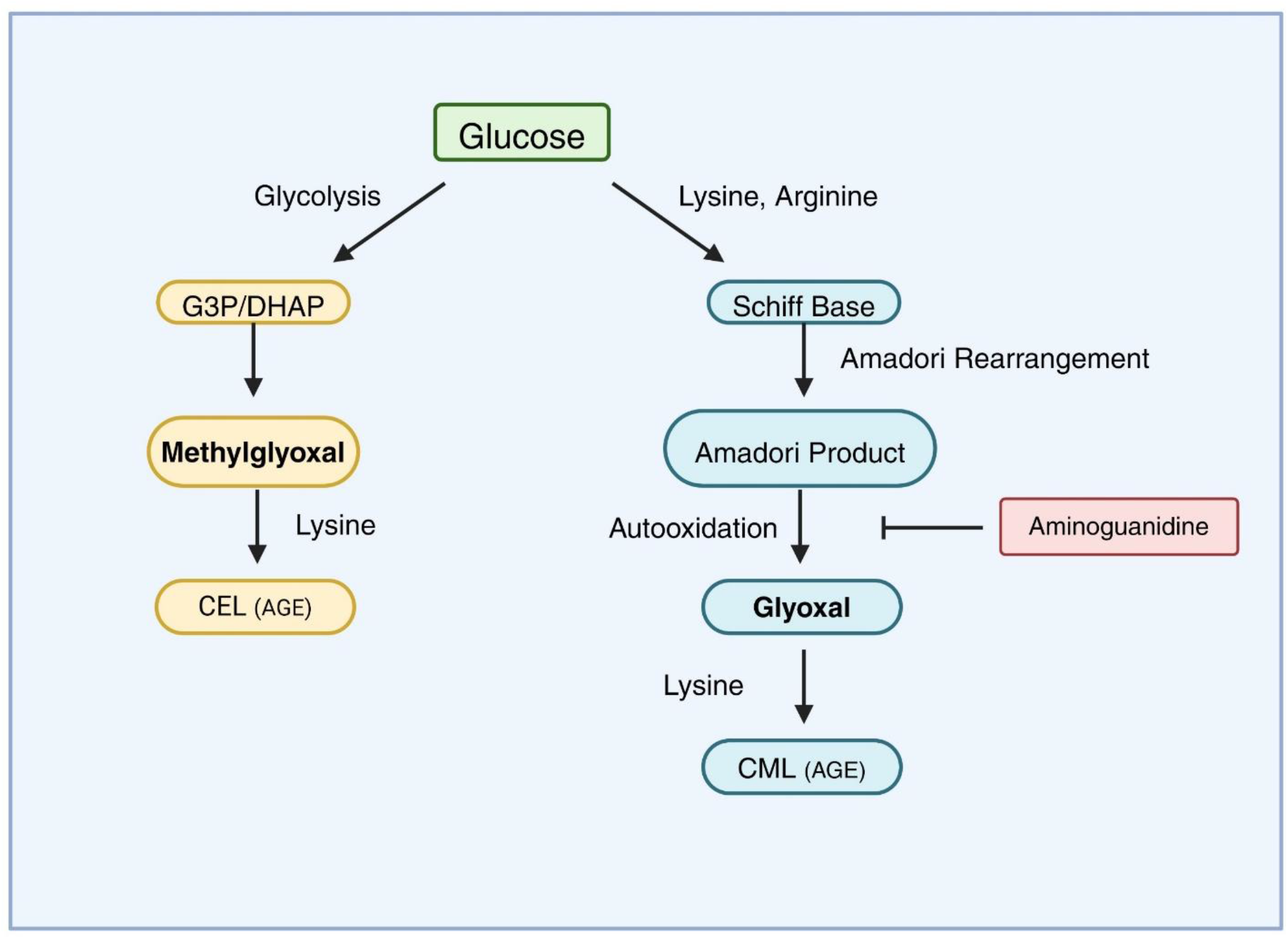
Formation of Methylglyoxal (MGO) and glyoxal (GO) from glucose. Glucose undergoes glycolysis to form glyceraldehyde-3-phosphate or Dihydroxyacetone phosphate and further oxidation steps produce methylglyoxal. The AGE Nε-(1-carboxyethyl) lysine or CEL is formed by methylglyoxal reaction with lysine residues on proteins. Glucose also forms a Schiff base through direct interaction with lysine and arginine. Through Amadori rearrangements of the Schiff base the Amadori product is formed which undergoes oxidation to yield glyoxal. Glyoxal can interact with lysine residues on proteins to yield the AGE Nε-(carboxymethyl)lysine (CML).

While our focus will be the endogenous production of MGO and GO it should also be noted that they can be produced by exogenous sources such as smoking or drinking. Due to the abundance of products and pathways involved in the formation of AGEs, the resulting damage that occurs from their accumulation is prevalent in many diseases such as but not limited to, cardiovascular, neurodegenerative, retinopathy, and kidney complications^5^. As shown briefly in figure 3, AGE-induced damage can occur through the binding of its receptor leading to oxidative stress, inflammation signaling, and cross-link formation resulting in impaired protein function and further cellular damage. ^6^

**Figure 3.**
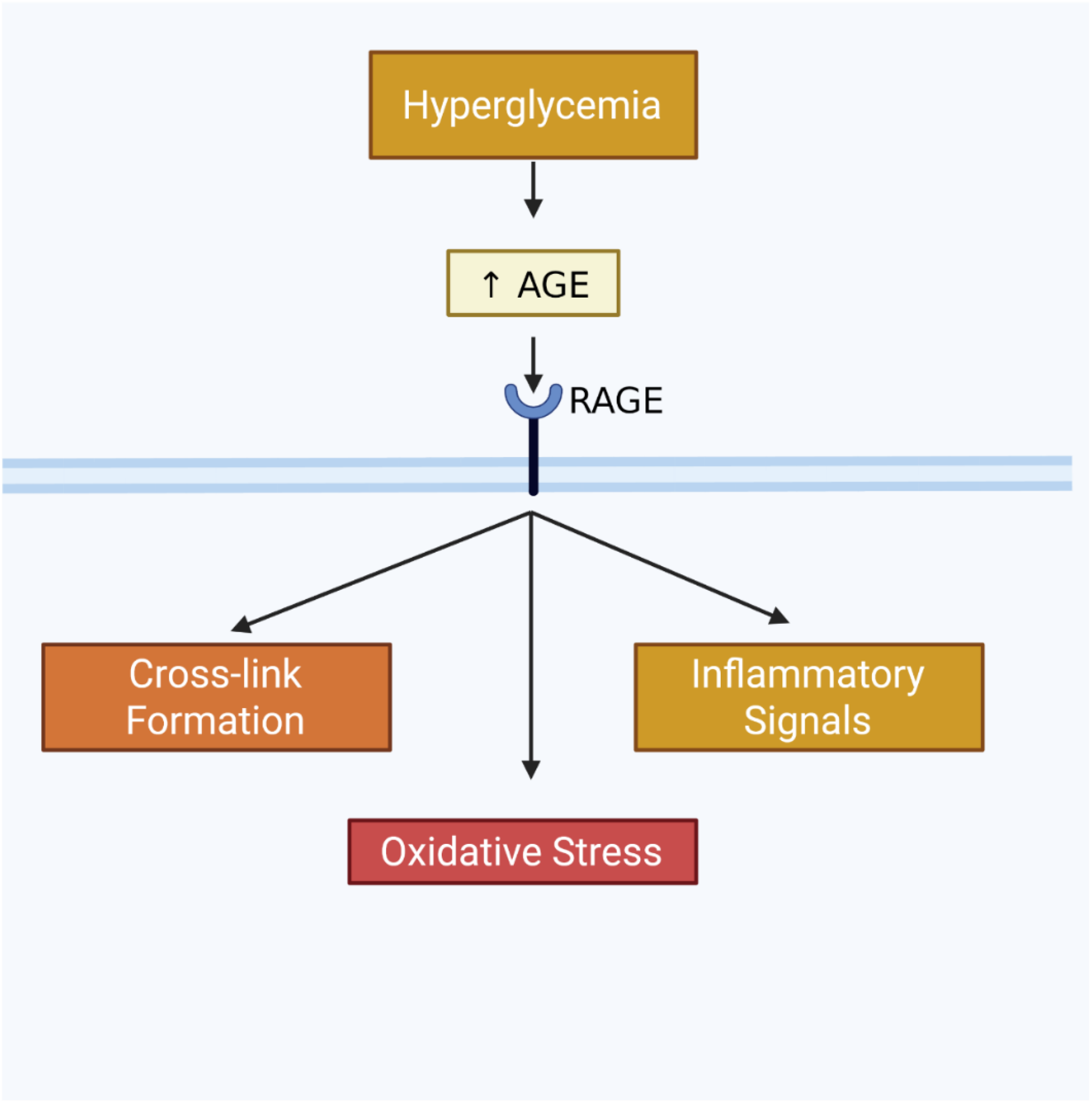
AGE-induced damage through RAGE Through interactions with its receptor RAGE, AGEs can cause oxidative stress, inflammation signaling, and cross-link formation.

However, while often overlooked, over half of diabetic patients experience significant musculoskeletal complications affecting their joints, tendons, muscles, and bones ^7^. Various musculoskeletal complications can arise due to advanced diabetes including sarcopenia, osteoporosis, and diabetic muscle infarction among others and greatly affects quality-of-life, morbidity rates, and higher incidence of cardiovascular complications in patients ^8,9^.

It is known that increased amounts of glucose in the blood stream during hyperglycemia can lead to increased accumulation of AGES through the formation of highly reactive fructose, which is later utilized in the Mailliard reaction, increasing the concentrations of AGES. At low levels, concentrations of AGES pose little to no threat. However, when concentrations exceed normal limits during hyperglycemic conditions, oxidative stress and tissue damage can occur due via the binding of AGEs to their receptor, RAGE. AGEs and their receptors have been extensively studied in the field, however, there is limited literature surrounding the direct effect of their dicarbonyl precursors. Most studies identified focus around the direct or downstream effects of AGEs on diabetic skeletal muscle. Furthermore, the studies that are based around the direct effect of these glycolaldehydes are modeled in non-skeletal muscle systems with various articles showing their effect in vascular smooth muscle cells, coronary endothelial cells, vascular endothelial cells, and bladder tissue to name a few^10–12^. Moreover, the aforementioned articles only focus on the metabolite Methylglyoxal with lacking data surrounding the other types of AGE precursors including Glyoxal. One article investigated the direct effect of Methylglyoxal in C2C12s, identifying MGO muscle damage through mitochondrial injury resulting in ATP depletion, decreased cell viability, and changes in fatty acid synthesis^13^. However, the study covered solely Methylglyoxal and explored the observed inflammatory aspect through a cytokine perspective rather than our proposed lipidomic lens. Data is also limited surrounding Glyoxal in most regards but especially in terms of its direct effect on a skeletal muscle model system, so more research is required to substantiate future claims.

Despite its prevalence, the mechanism of action of diabetic skeletal muscle pathology is poorly understood. As such, the objective of this study is to investigate the hypothesis: Glucose-derived glycolaldehydes, glyoxal and methylglyoxal formed in hyperglycemia cause damage to the skeletal muscle myoblasts through oxidative stress and formation of pro-inflammatory eicosanoids upon activation of phospholipase A_2_ (PLA_2_) and cyclooxygenase (COX).

The work of this project aims to fill these gaps in knowledge by exploring dicarbonyl-induced damage in a previously unexplored skeletal muscle model system, and to identify a novel lipidomic/oxidative stress mechanism by which this damage occurs. This project will also aim to propose various therapeutic targets to attenuate observed damage through the identified pathway.

## Materials & Methods

Skeletal muscle mouse myoblasts (C2C12s) were obtained from the American Type Culture Collection (ATCC;1-6 passages) and grown in Dulbecco’s Modified Eagle Medium (DMEM) containing 4.5g/l D-Glucose, L-glutamine, absence of sodium pyruvate, 10% fetal bovine serum, 1% Antimycotic-Antibiotic, and 1% Penicillin-streptomycin. Fetal bovine serum (FBS), minimum essential medium (MEM), phosphate buffer saline (PBS) in the absence of calcium and magnesium chloride, and TrypLE were obtained from Gibco Invitrogen Corp. [^3^H]-Thymidine was obtained from the American Radiolabeled Chemicals, Inc. Lactic acid dehydrogenase cytotoxicity assay (LDH) kit, and (3-(4,5-dimethylthiazol-2-yl)-2,5-diphenyl tetrazolium bromide) (MTT) were obtained from the Sigma Chemical Co. The cPLA_2_ Activity Assay kit was obtained from Cayman Chemical Company. 6-carboxy-Alamar Blue reagent and 2,7-dichlorodihydroxyfluorescein diacetate dicarboxy methyl ester (DCFDA) was purchased from ThermoFisher Scientific. Glyoxal trimer dihydrate and Methylglyoxal for cell treatment was obtained from Sigma-Aldritch.

### Cell Culture

The C2C12 mouse myoblast cells (ATCC; 1-6 passages) were cultured in DMEM supplemented with 10% FBS, 1% Antimycotic-Antibiotic and 1% Penicillin-streptomycin. Cells were cultured up to 90–100% confluence, lifted using TrypLE, and subcultured in sterile 35-mm or 60-mm dishes and in 96-well plates under a sterile and humidified atmosphere of 95% air–5% CO2 at 37 °C ^14^.

### Cell Morphology

Morphological changes in C2C12s grown in 35-mm sterile dishes up to 90% confluence, following their exposure to different concentrations of glyoxal (0 −2 mM) in absence or presence of chosen pharmacological agents (0–5.0 mM) in MEM for 0–24 h at 37 °C in a humidified environment of 5% CO2 – 95% air, were examined. Images of cell morphology were digitally captured with the Olympus 1 × 50 microscope at 10 X and 20 X magnification^15^.

### Cytotoxicity Assays

#### LDH

Cytotoxicity was assayed by the release of Lactic Acid Dehydrogenase (LDH). C2C12s were grown up to 70% confluence in 3.5 cm^2^ dishes (12-well culture plates) and exposed to various glyoxal concentrations (0-2 mM) in the presence or absence of antioxidant compounds (NAC or GSH), or Metformin (0-10 mM) for 24-48 h. Following incubation, the supernatant was removed and release of LDH was determined via spectrophotometrically methods and expressed at absorbance differences (490 −690 nm) ^15^.

#### Reduction of MTT

C2C12s were grown up to 70% confluence in 3.5 cm^2^ dishes (12-well culture plates) and exposed to various glyoxal concentrations (0-2 mM) in the presence or absence of antioxidant compounds (NAC), Inflammation and PLA_2_ inhibitors (AACOCF3, Ibuprofen,), or Metformin (0-10 mM) for 24-48 h. Following incubation, MTT solution was added and plates were additionally incubated for 2-3 h. The solution was removed; MTT solubilizing solution was added to each well equivalent to the original culture volume and absorbance was read. MTT reduction was expressed as the difference between 570nm and 690 nm.

#### Trypan Blue uptake

C2C12s were grown to 70% confluence in 6-well culture plates in complete medium and exposed to various glyoxal concentrations (0-2 mM) for 24h. Following incubation, treatment media was removed, 0.2% of Trypan Blue Solution diluted in PBS was added to each well and incubated for 40mins. The solution was then removed, washed gently twice with PBS and lifted using 1% Triton X-100. Trypan Blue uptake was measured spectrophotometrically in a 96-well plate at 570 nm utilizing an Agilent plate reader. Data was expressed as absorbance in nanometers.

#### ATP Luminescence

C2C12s were grown to 50-60% confluence in 6-well culture plates and treated with Glyoxal at 1 mM in DMEM, 5 mM Metformin in DMEM, DMEM Alone, and 1 mM glyoxal and 5 mM Metformin co-treatment in DMEM in triplicate. Following a 24-hour incubation period, cell lysates were prepared as samples. ATP standards and experimental standards were run in accordance with the ATP Determination Kit from ThermoFisher Scientific. ATP levels of the samples were determined from the sample curve and expressed as luminescence.

### [3H]-Thymidine Incorporation Assay for Cell Proliferation

C2C12 cells were grown to 60% confluence in 35-mm dishes. Complete medium was removed from the culture dishes, and treatment solutions were added to the dishes. The treatment medium was then removed and 1 mL of [3H]-thymidine (1.0 μCi/mL) in DMEM was added to each well and incubated for 24 h. After incubations, the [3H] radioactivity was counted in the Packard Tri-carb 2900TR Liquid Scintillation Counter (Gurney et al., 2022).^15^ Cell replication was expressed as DPM of [3H]-thymidine incorporation into cells/dish.

### ROS Measurements

Formation of cellular ROS in the C2C12 cells and by the mitochondria was measured by the fluorescence of DCFDA by utilizing the fluorescence plate reader.

General ROS formation in 35-mm dishes was measured by 2′,7′-dichlorofluorescein (DCF) fluorescence in cells preloaded with 10 μM DCFDA for 30 min in complete MEM at 37 °C in a 5% CO2 – 95% air environment prior to exposure to glyoxal (0–2 mM) in absence or presence of antioxidants, inhibitors, or metformin for 3 or 6 hours. Following treatment incubation, media was read in a plate reader at 490 nm excitation and 530nm emission. The cells were then washed with PBS (2 x 1mL each), collected using a Teflon cell scraper in PBS, transferred to 1.5mL microcentrifuge tubes and spun down at 8000 x g for 10 minutes. The pellets were sonicated on ice at a probe setting of 2 for three 15-second intervals with 10-second breaks. The fluorescence of oxidized DCF in the lysates was then measured at the same excitation and emission wavelengths ^15^.

### Cellular Redox Measurement

C2C12s were cultured in 22-mm wells at 70% confluence in complete medium and exposed to various glyoxal concentrations (0-2 mM) for 24h. Following incubation, treatment media was removed, Alamar blue reagent was added and incubated for 2h. Absorbance signals were read at wave lengths 600 nm and 570 nm using the Agilent plate reader. Alamar Cellular redox was determined spectrophotometrically by the reduction of Alamar Blue ^16^.

### Mitochondrial Cardiolipin Determination

C2C12s were cultured in 35-mm wells at 70% confluence in complete medium and exposed to various glyoxal concentrations (0-2 mM) for 24h in the absence or presence of Metformin. Cells were stained with 10-nonyl acridine orange (10-NAO) at a concentration of 500nm in subdued light for 40 mins. Following incubation cells were either imaged using the Olympus 1 × 50 fluorescent microscope at 20X magnification or read via prepared cell lysates. Cell lysates were prepared following incubation by washing twice with 1mL of PBS, scraping adherent cells with a Teflon scraper, collecting in a 1.5 mL centrifuge tube, and sonicating for 3 15-second intervals with 10 second breaks in between. Lysates were then plated in a 96-well plate and cardiolipin content of mitochondria was determined fluorometrically on a plate reader. Microscopy results were expressed as mean fluorescence in fold change and lysates were expressed as Fluorescence 485/20,528/20.

### [^3^H]-Arachidonic Acid Release Assay

Cells were grown to 60-70% confluence in 6-well culture plates and treated with Glyoxal concentrations of 0.1, 1, and 2mM. PLA_2_ activity was assayed by the release of [^3^H]-Arachidonic Acid from cells following treatment of cells with glyoxal according to established protocol^17^.

### cPLA_2_ Activity Assay

Cells were grown to 70-80% confluence in 6-well culture plates and treated with 1 mM Glyoxal for 15-, 30-, and 60-minute time points. Cell lysates were prepared and a BSA Assay was run to determine protein concentration. Protein content between samples was normalized and the cPLA_2_ Activity Assay was performed in accordance with the Cayman Chemical cPLA_2_ Assay Kit^18^.

### Eicosanoid Analysis

Eicosanoids formed in cells following treatment with glyoxal were analyzed by the LC-MS method^19–20^.

### Fluorescent Imaging

Cells were cultured in a 96-well culture plate and treated with 1 mM glyoxal and 5mM metformin at 70-80% confluency. Following treatment, cells were washed twice with ice-cold PBS and fixed with 4% paraformaldehyde in PBS for 15-20 mins at room temperature. Cells were then washed twice with cold PBS and permeabilized with 0.25% Triton X-100 in PBS containing 0.01% Tween-20 (PBST) for 5 minutes at room temperature. Cells were washed twice with PBST and blocked with 1% bovine serum albumin (BSA) in PBST for 15-20 minutes at room temperature.

#### cPLA_2_ Activity Staining

Following blocking, cells were incubated with anti-cPLA_2_ primary antibody (Santa Cruz Biotechnology, catalog no. sc-438; 1:25 dilution in blocking buffer) overnight at 4°C. Cells were washed three times with PBST before incubation with the secondary antibody Goat anti-Mouse IgG (H+L) Highly Cross-Adsorbed Secondary Antibody, Alexa Fluor™ Plus 555 (catalog no. A10042; 1:100 dilution in blocking buffer) for 2h at room temperature. Cells were washed five times with PBST and counterstained with DAPI (100 ng/mL in PBS) for 5 min at room temperature. Cells were washed three times with PBS and fluorescently imaged immediately at 20X magnification. Values were averaged and plotted and standard deviations added using Fiji and ImageJ.

#### Phalloidin Staining

Following blocking as described previously, cells were incubated with phalloidin (1:100 dilution in blocking buffer) for 2 h at room temperature. Cells were washed five times with PBST and counterstained with DAPI (10ng/mL in PBS) for 5 min at room temperature. Cells were washed with PBS and imaged immediately at 20X magnification. Values were averaged and plotted and standard deviations added using Fiji and ImageJ.

#### Statistical Analysis of Data

Values are presented as the means of triplicate sample measurements. Data were analyzed using on-way ANOVA, followed by Dunnett’s post hoc test for comparisons against the control group. Dose-response and inhibition experiment planned comparisons were performed in R using estimated marginal means, with Holm adjustment for multiple comparisons. Statistical significance was set at p <0.05.

## Results

### Glycolaldehydes Inhibit Cellular Proliferation in Skeletal Muscle

Few studies have shown the direct impact of glyoxal and methylglyoxal on skeletal muscle cells. Here we investigated the effect on cellular proliferation in response to increasing doses of our glycolaldehydes. Figure 4 measured cellular proliferation as the extent of [3^H^]-Thymidine incorporation in intact cells following treatment with Glyoxal 0.5 mM-2 mM.

**Figure 4.**
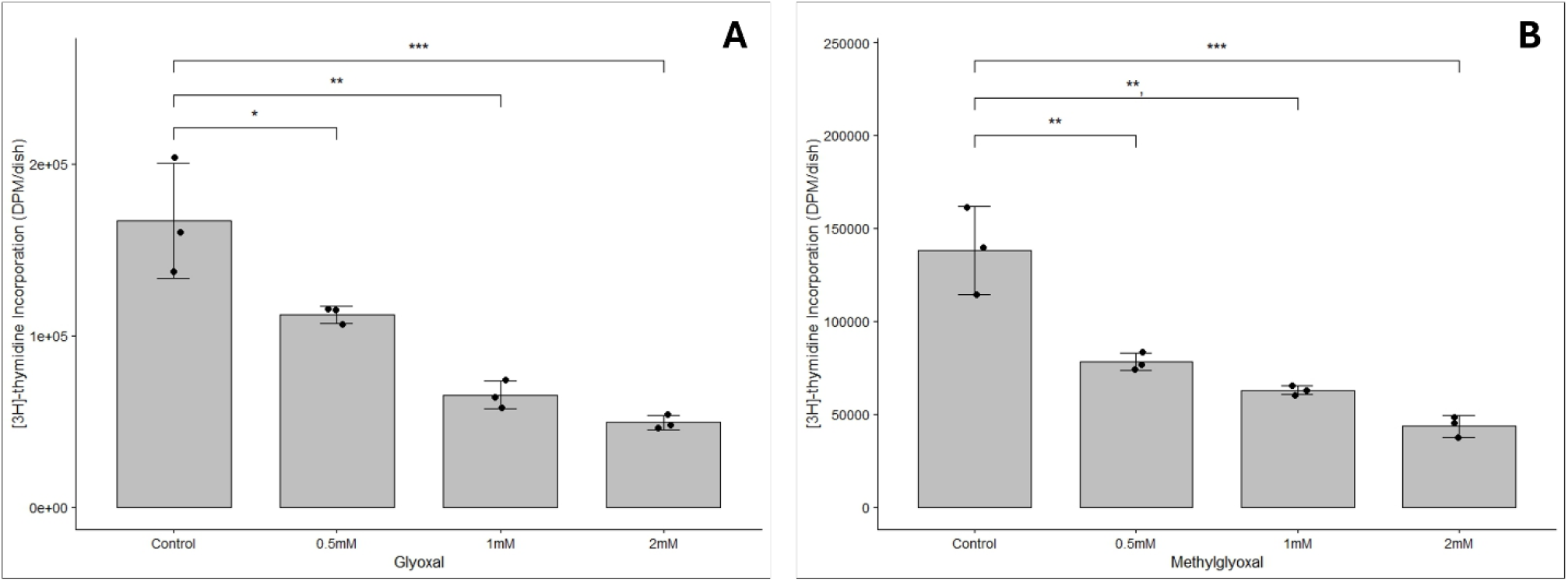
Glyoxal and Methylglyoxal inhibit cellular proliferation in a dose-dependent fashion. C2C12s (60% confluency/ 35-mm dish) were treated with **(A)** glyoxal (0.5mM, 1mM, 2mM) or **(B)** methylglyoxal (0.5mM, 1mM, 2mM) for 24 h. Following treatment, cells were labeled with [3H]-thymidine (1.0µCi/mL in DMEM) for 24 h. Data represents means ± SD. Compared to untreated cells, * = significance at p <0.05, ** = significance at p < 0.01, *** = significance at p<0.001.

All concentrations of glyoxal 0.5-2 mM showed a dose-dependent statistically significant decrease in cellular proliferation following 24 hours of treatment with p < 0.05. A 32%, 60%, and 70% decrease from the control was observed for the 0.5 mM, 1 mM, and 2 mM doses respectively. A similar trend at significance p < 0.05 was observed in methylglyoxal at doses 0.5 mM, 1 mM, and 2 mM with a 43.4%, 54.4%, and 68.5% decrease respectively.

### Glycolaldehydes Induce Cytoskeletal Damage and Membrane Tears

To further investigate how MGO and GO induce damage in C2C12s, membrane integrity assays Trypan Blue and LDH were run at the concentrations of 0.5mM, 1mM, and 2 mM for MGO and GO. Figure 5A demonstrates glyoxal uptake of trypan blue at a 1.5-fold increase at the highest dose compared to control but the result remained statistically insignificant. However, Figure 5C shows the extent of lactate dehydrogenase release at a dose-dependent increase of up to 3-fold compared to controls at p < 0.05. Similarly, MGO significantly demonstrated up to a 1.3-fold increase compared to control for trypan blue uptake in Figure 5B and up to a 1.7-fold increase in the release of LDH in Figure 4D.

**Figure 5.**
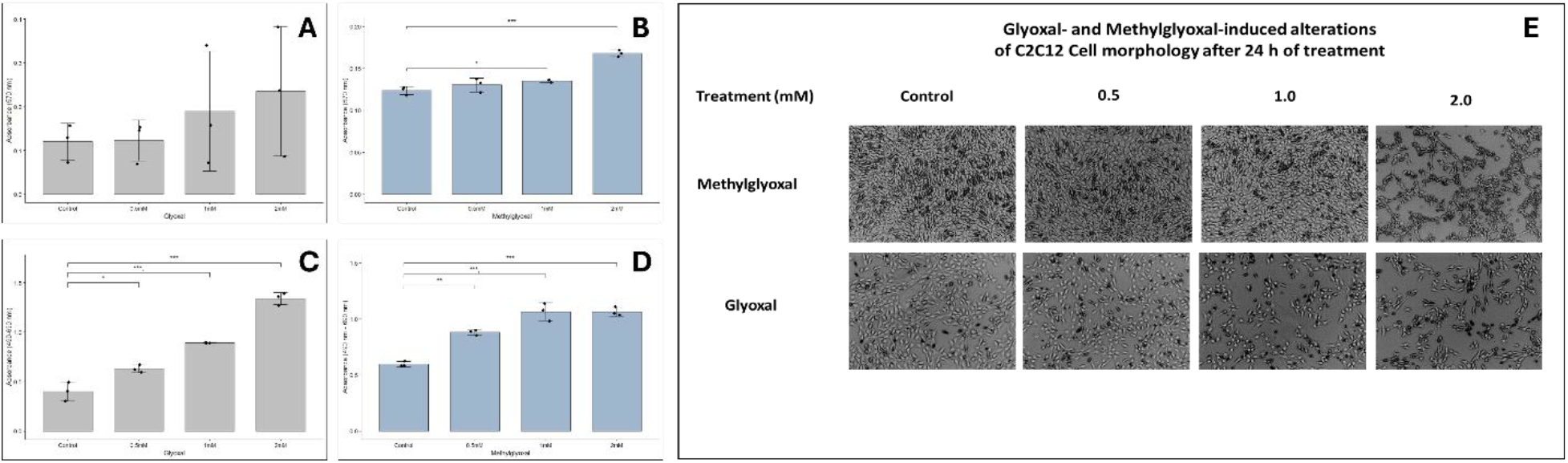
MGO and GO induce cytoskeletal damage and cell membrane tears. C2C12s (70% confluency in 6-well or 12-well culture plates) were treated with **(A, C)** glyoxal (0.5mM, 1mM, 2mM0 or **(B, D)** methylglyoxal (0.5mM, 1mM, 2mM) for 24h. Following Incubation uptake of trypan blue **(A-B),** release of LDH **(C-D),** and brightfield morphology images at 10X magnification **(E)** were measured as described in Materials and Methods. Data represents means ± SD of three independent experiments. Compared to untreated cells, * = significance at p <0.05, ** = significance at p < 0.01, *** = significance at p<0.001.

Additionally, Figure 5E shows a visualization of cellular morphological changes for increasing doses of MGO and GO where cell shape changes from elongated at baseline to rounded and compromised at 2mM.

### Glyoxal Alters Metabolic and Redox Characteristics and Increases ROS Production

Previous findings from our lab have established the presence of metabolic and redox changes within the model system of BPAECs. ^21^ Similar studies were conducted within our model system. Figure 6A shows a decrease in absorbance values in the MTT assay of up to 98.8% at 2 mM compared to controls at p < 0.01. Figures 6B and 6C show similarly significant findings for methylglyoxal with a 28.1% decrease at 2 mM and a 98.5% decrease at 3 mM.

**Figure 6.**
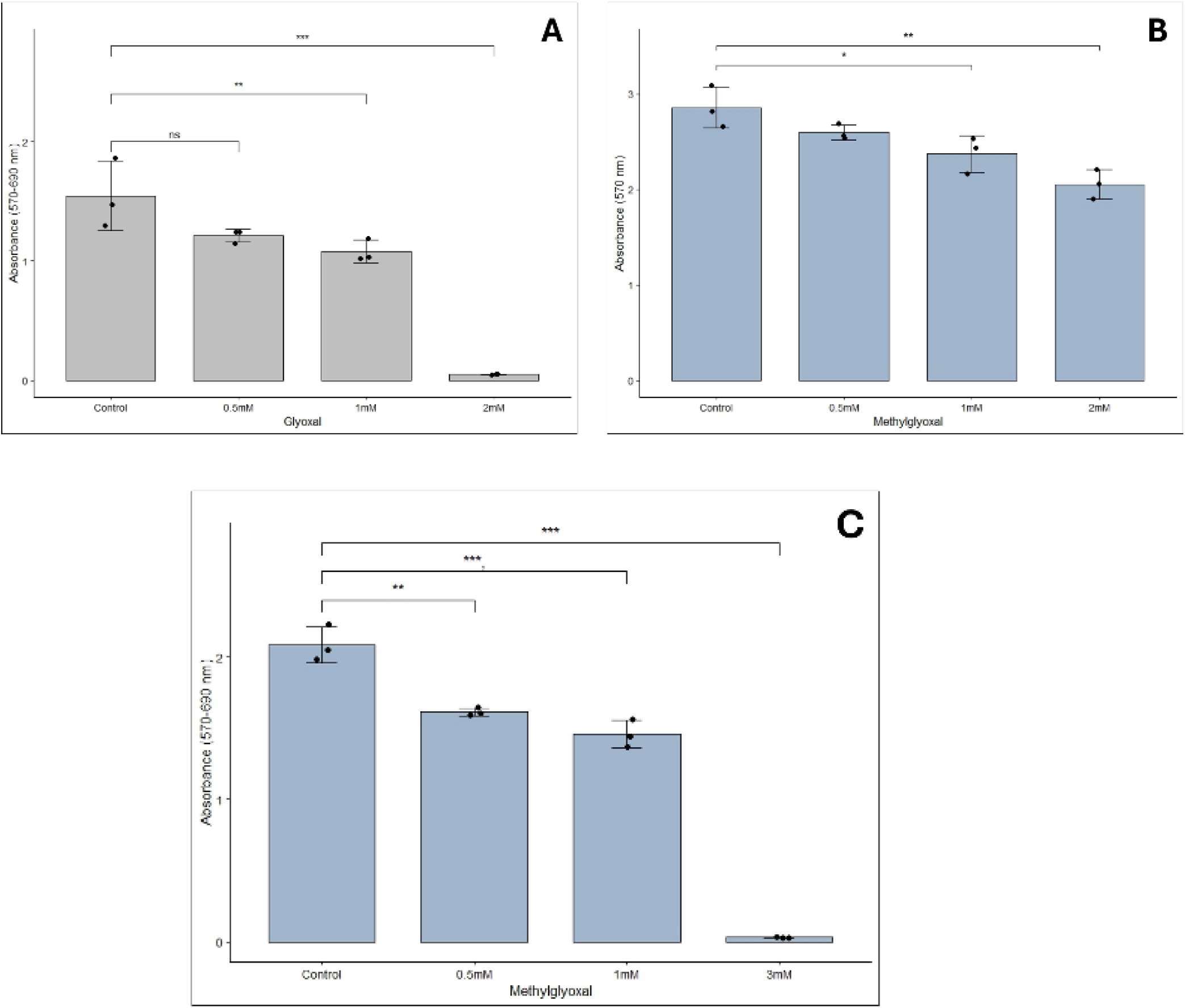
MGO and GO alter metabolic activity in a dose-dependent manner. C2C12s (70% confluency in 12-well culture plates) were treated with **(A)** glyoxal (0.5mM, 1mM, 2mM), **(B)** methylglyoxal (0.5mM, 1mM, 2mM), or **(C)** methylglyoxal (0.5mM, 1mM, 3mM) for 24 h. Following treatment, cellular metabolic activity was assessed by MTT reduction as described in Materials and Methods. Data represent means ± SD of triplicate sample measurements. Compared to untreated cells, * = significance at p < 0.05, ** = significance at p < 0.01, *** = significance at p < 0.001.

The redox status of the cells was also significantly changed in both treatments with glyoxal having up to a 1.8-fold increase in redox inhibition (Figure 7A), and methylglyoxal up to a 3.2-fold increase in inhibition (Figure 7B) at p < 0.001.

**Figure 7.**
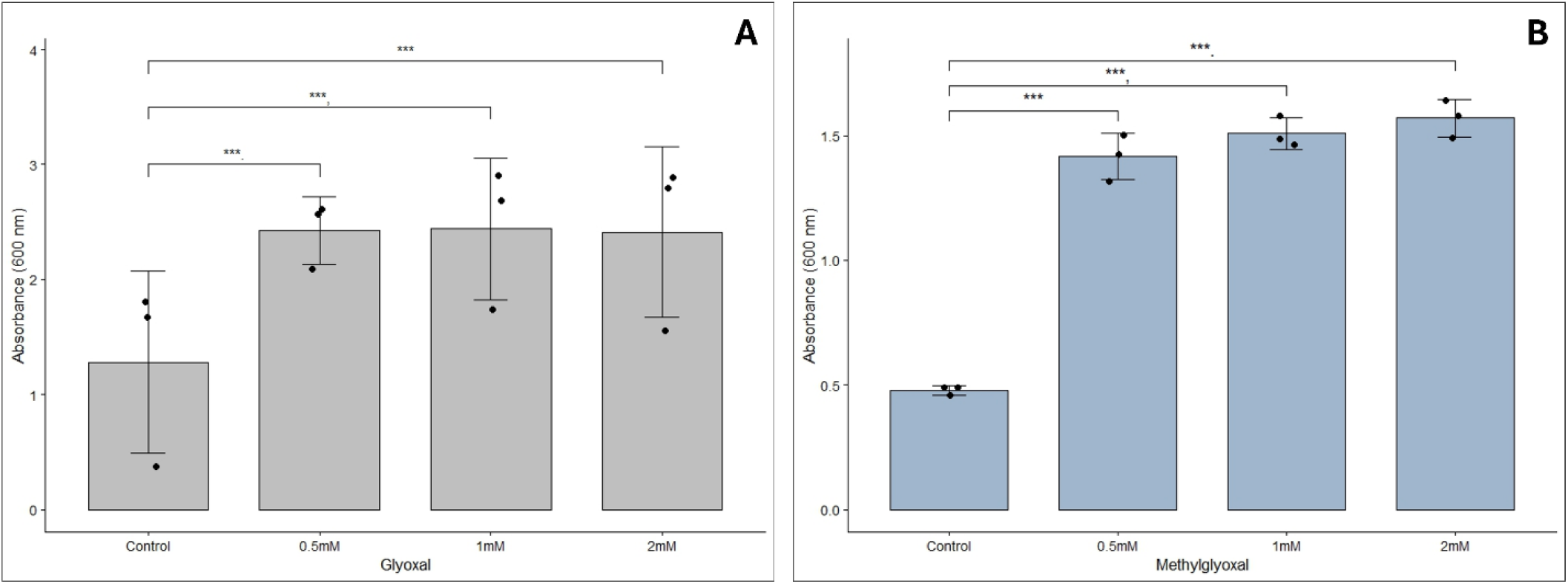
GO and MGO decrease the redox status of skeletal muscle cells in a dose-dependent fashion. C2C12s (70% confluency in culture wells) were treated with **(A)** glyoxal or **(B)** methylglyoxal at 0.5mM, 1mM, and 2mM for 24 h. Following treatment, cellular redox status was measured using Alamar Blue reagent after a 2 h incubation, as described in Materials and Methods. Data represent means ± SD of triplicate sample measurements. Compared to untreated cells, * = significance at p < 0.05, ** = significance at p < 0.01, *** = significance at p < 0.001.

Furthermore, the levels of ATP produced were measured (Figure 8) in glyoxal 1 mM, metformin 5 mM, and a co-treatment. Results were statistically significant with an 85.6% decrease in ATP production at glyoxal 1 mM compared to controls, and a 3.7-fold increase in ATP production during co-treatment compared to glyoxal alone. The co-treatment group, however, did demonstrate a 46.5% decrease in ATP compared to controls.

**Figure 8.**
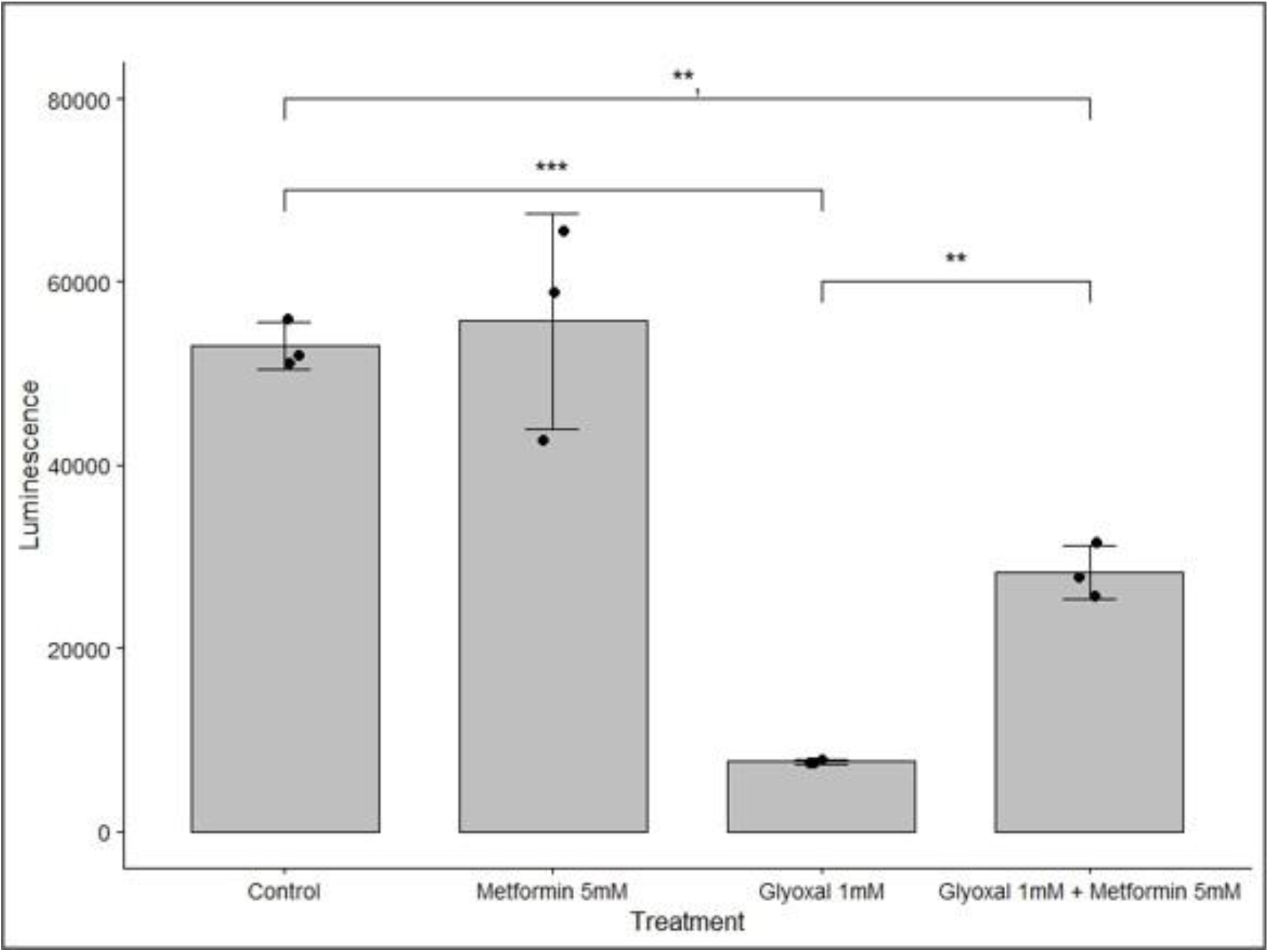
Glyoxal inhibits the production of ATP which is significantly attenuated by metformin. C2C12s (50–60% confluency in 6-well culture plates) were treated with DMEM alone, glyoxal (1 mM), metformin (5 mM), or glyoxal (1 mM) and metformin (5 mM) in combination for 24 h. Following treatment, cell lysates were prepared and ATP levels were determined using an ATP determination kit. ATP levels were calculated from the standard curve and expressed as luminescence. Data represent means ± SD of triplicate samples. Compared to untreated cells, * = significance at p < 0.05, ** = significance at p < 0.01, *** = significance at p < 0.001.

Finally, Figure 9 demonstrates a significant increase in ROS production in methylglyoxal at a 1.6-fold increase compared to controls for all doses.

**Figure 9.**
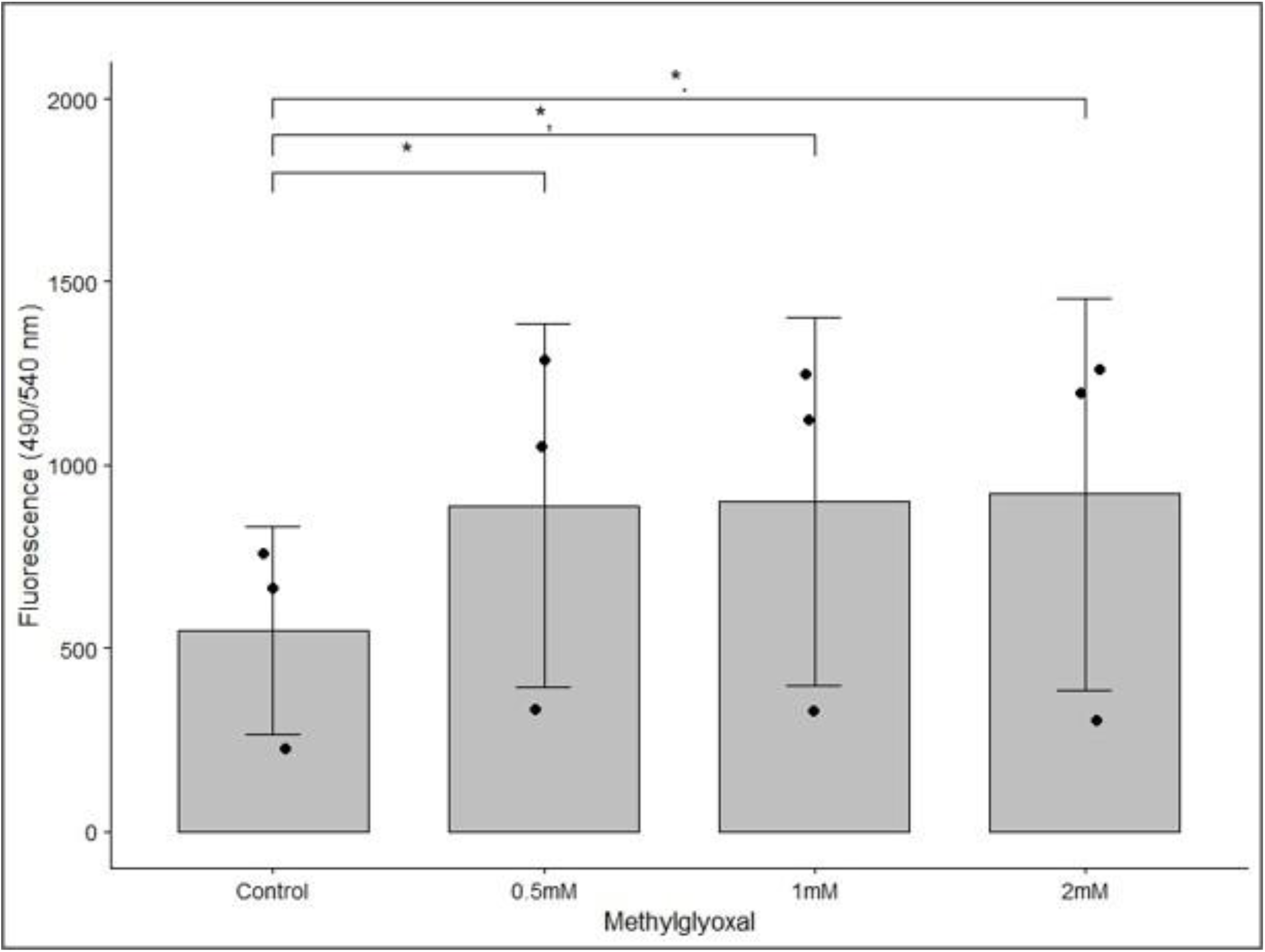
MGO increases the production of ROS in skeletal muscle. C2C12s were loaded with DCFDA (10 μM) for 30 min before treatment with methylglyoxal 0.5mM, 1mM, and 2mM. Following treatment, ROS production was assessed by measuring DCF fluorescence at 490 nm excitation and 530 nm emission, as described in Materials and Methods. Data represent means ± SD of triplicate sample measurements. Compared to untreated cells, * = significance at p < 0.05, ** = significance at p < 0.01, *** = significance at p < 0.001.

#### PLA_2_ is Activated by Glyoxal

Due to observed cytoskeletal changes, phospholipase A2 activity was measured to observe a possible inflammatory aspect in the mechanism. Figure 10A shows a significant increase in the amount of [^3^H]-Arachidonic Acid release at 0.5 mM and 1 mM of up to 1.3-fold compared to controls. Similarly, glyoxal demonstrated the time point for peak PLA_2_ activation at 30 minutes with a drop in activity at 60 minutes. This result is consistent with current literature in other model systems suggesting a rapid activation of inflammatory pathways.^10^

**Figure 10.**
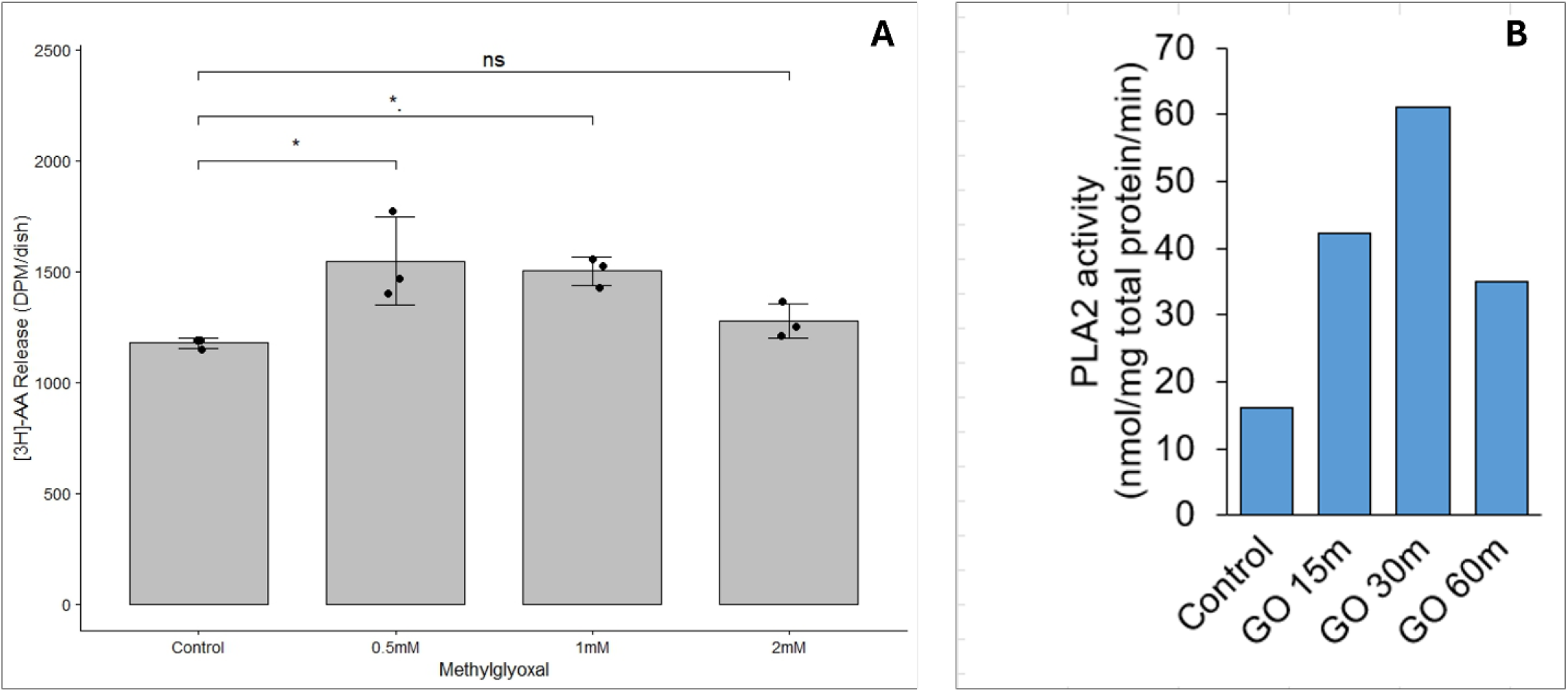
MGO and GO induce the activation of PLA_2_. C2C12s (60–70% confluency in 6-well culture plates) were treated with glyoxal at the indicated concentrations, and PLA_2_ activity was assessed by measuring the release of [³H]-arachidonic acid (AA) from cells **(A)**. In a separate assay, C2C12s (70–80% confluency in 6-well culture plates) were treated with glyoxal (1 mM) for 15, 30, or 60 min. Cell lysates were prepared, protein concentrations were determined, and cPLA_2_ activity was measured using a commercial assay kit **(B)**, as described in Materials and Methods. Data represent means ± SD of triplicate sample measurements. Compared to untreated cells or the corresponding control group, * = significance at p < 0.05, ** = significance at p < 0.01, *** = significance at p < 0.001.

### Key Inflammatory Markers are Upregulated

As a result of cPLA_2_ activation, figures 11A-H demonstrates the dose dependent increase of inflammatory markers PGE1, PGE2, 15-keto PGE2, PGA2, PGD2, Δ12-PGJ2, PGF2α, and 6-keto PGF1α produced from the release of arachidonic acid and subsequent activation of COX and LOX

**Figure 11.**
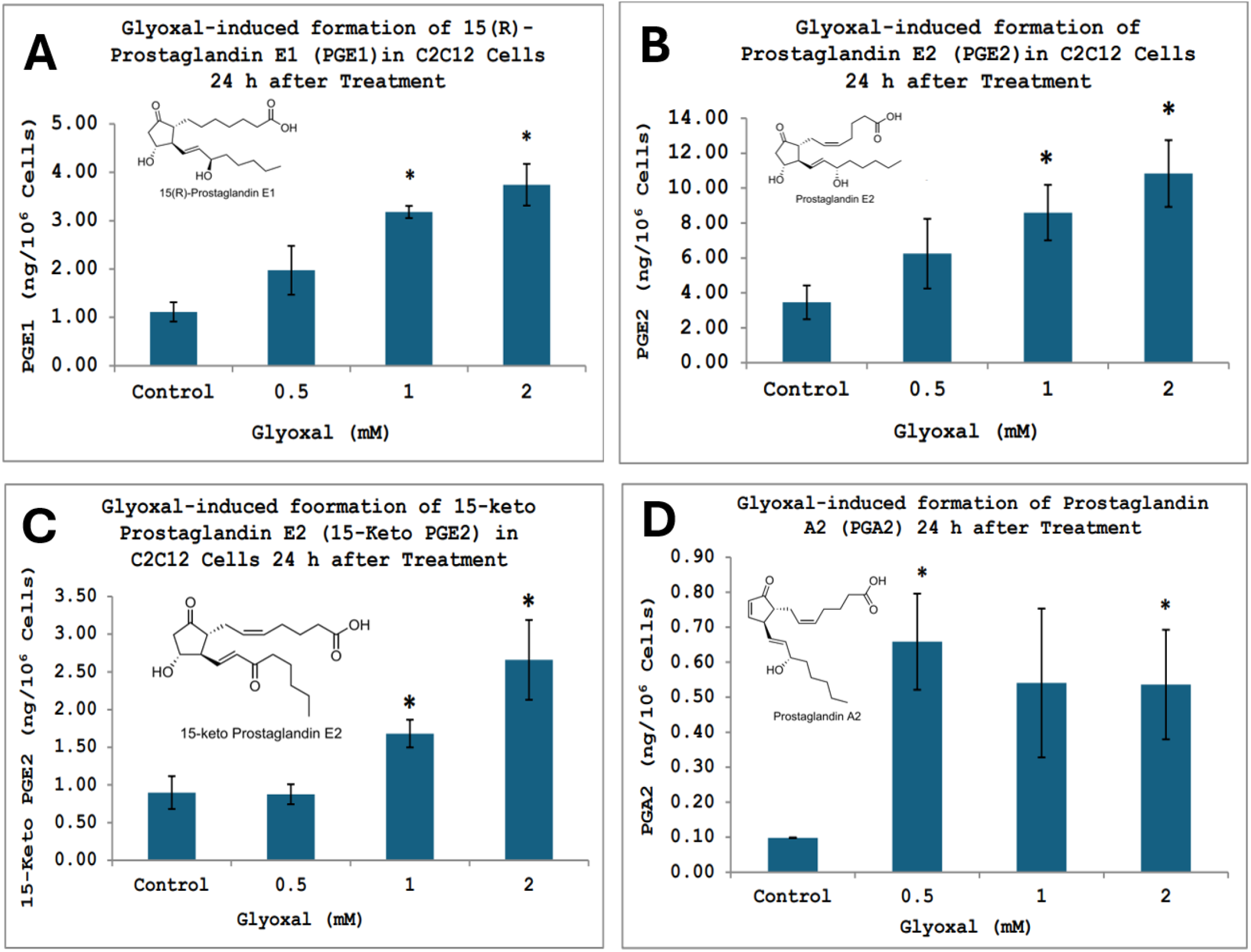
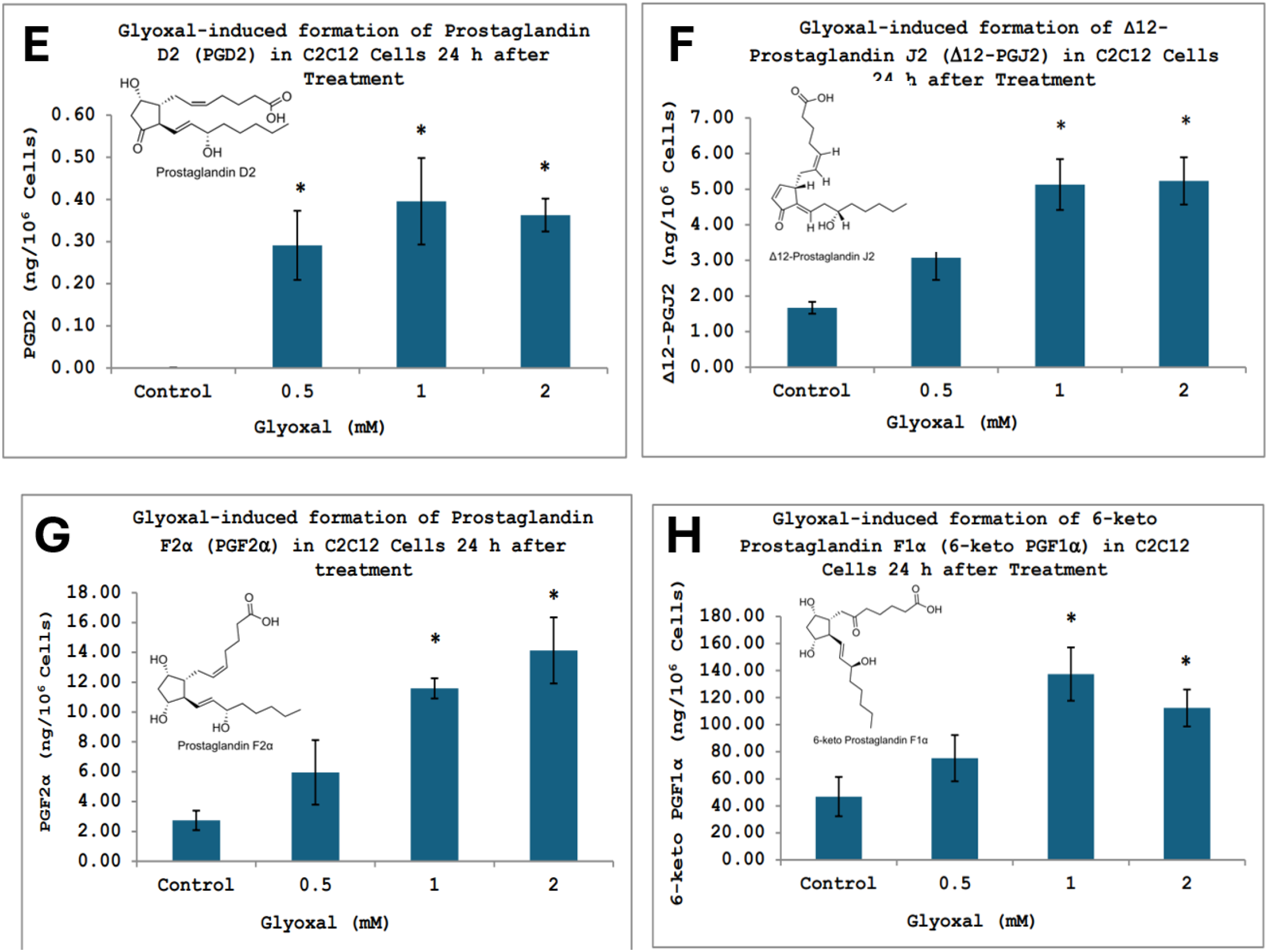
Glyoxal increases the production of key inflammatory prostaglandins. C2C12s were treated with glyoxal under the indicated experimental conditions. Following treatment, cellular eicosanoids were analyzed by liquid chromatography–mass spectrometry (LC-MS), as described in Materials and Methods. The levels of the indicated inflammatory prostaglandins and related eicosanoids were measured across the eight experimental groups. Data represent means ± SD of triplicate sample measurements. Compared to the corresponding control group, * = significance at p < 0.05, ** = significance at p < 0.01, *** = significance at p < 0.001.

Figure 11A and 11B show 2.8- and 3.3-fold increase in PGE1 at 1mM and 2mM, and a 2.4- and 3.1-fold increase in PGE2 at 1mM and 2mM. Figures 11C and 11D demonstrate significant increases in 15-keto PGE2 and PGA2. 15-keto PGE2 showed a 1.8- and 2.9-fold increase at doses 1 mM and 2mM, and PGA2 showed a 6.6-fold increase at 0.5mM, no significance at 1mM, and a 5.4 increase at 2mM. Figures 11E and 11F demonstrate increases in PGD2 and Δ12-PGJ2. PGD2 showed statistically significant increases at all doses with non-detectable levels at baseline. Δ12-PGJ2 demonstrated significance at 1 mM and 2 mM with 3.0- and 3.1-fold increases respectively.

Finally, figures 11G and 11H show increases in PGF2α and 6-keto PGF1α. PGF2α demonstrated a 4.2- and 5.1-fold increase at doses 1 mM and 2 mM, whereas 6-keto PGF1α demonstrated a 2.9- and 2.3-fold increase at the same doses.

### Loss of Mitochondrial Membrane Lipid

In response to observed mitochondrial dysfunction, loss of cardiolipin, a lipid within the inner matrix of the mitochondrial membrane, was measured at varying concentrations of glyoxal (0.5-2 mM). Figure 12A shows up to a 73.7% loss of cardiolipin at 1mM compared to controls with slightly less loss (60.5%) at the highest dose. Figure 12B, however, shows a clear statistically significant decrease in the amount of fluorescently labeled cardiolipin through NAO staining, with the highest loss of lipid observed at the 2 mM dose.

**Figure 12.**
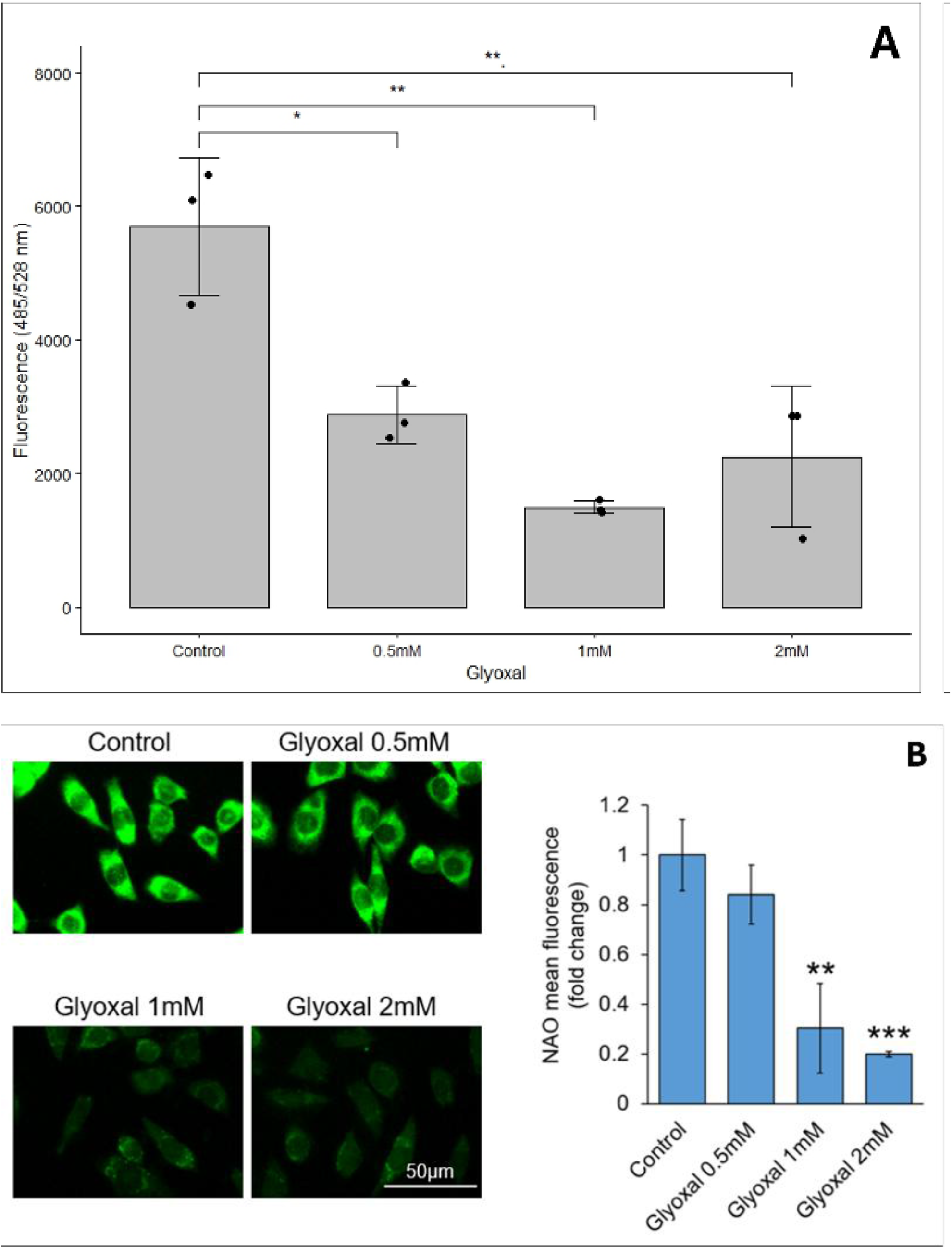
Glyoxal causes the loss of cardiolipin from mitochondrial membranes. C2C12s (70% confluency in culture wells) were treated with glyoxal at the indicated concentrations for 24 h in the absence or presence of metformin. Following treatment, cells were stained with 10-nonyl acridine orange (10-NAO; 500 nM) for 40 min. Mitochondrial cardiolipin was assessed by fluorometric measurement of cell lysates **(A)**, and by fluorescence microscopy at 20X magnification **(B)** as described in Materials and Methods. Microscopy results **(C)** are expressed as fold change in mean fluorescence, and lysate measurements are expressed as fluorescence intensity. Data represent means ± SD of triplicate sample measurements. Compared to untreated cells, * = significance at p < 0.05, ** = significance at p < 0.01, *** = significance at p < 0.001.

#### NAC Protect from Metabolism loss in Moderation

To establish therapeutic action points within the pathway, cells were treated with the antioxidant N-acetylcysteine (NAC; 0.25-1 mM) in the absence or presence of glyoxal 1 mM and redox and mitochondrial function were evaluated. Figure 13A demonstrates limited NAC amelioration of glyoxal-induced redox changes resulting in only a 7.9% decrease in redox loss at the highest dose compared to glyoxal alone, and up to a 1.41-fold increase in redox loss at 1 mM NAC alone compared to controls. Significant attenuation, however, was observed in a measurement of mitochondrial functionality (Figure 13B) with up to a 2.3-fold increase in metabolic function compared to glyoxal at 1 mM.

**Figure 13.**
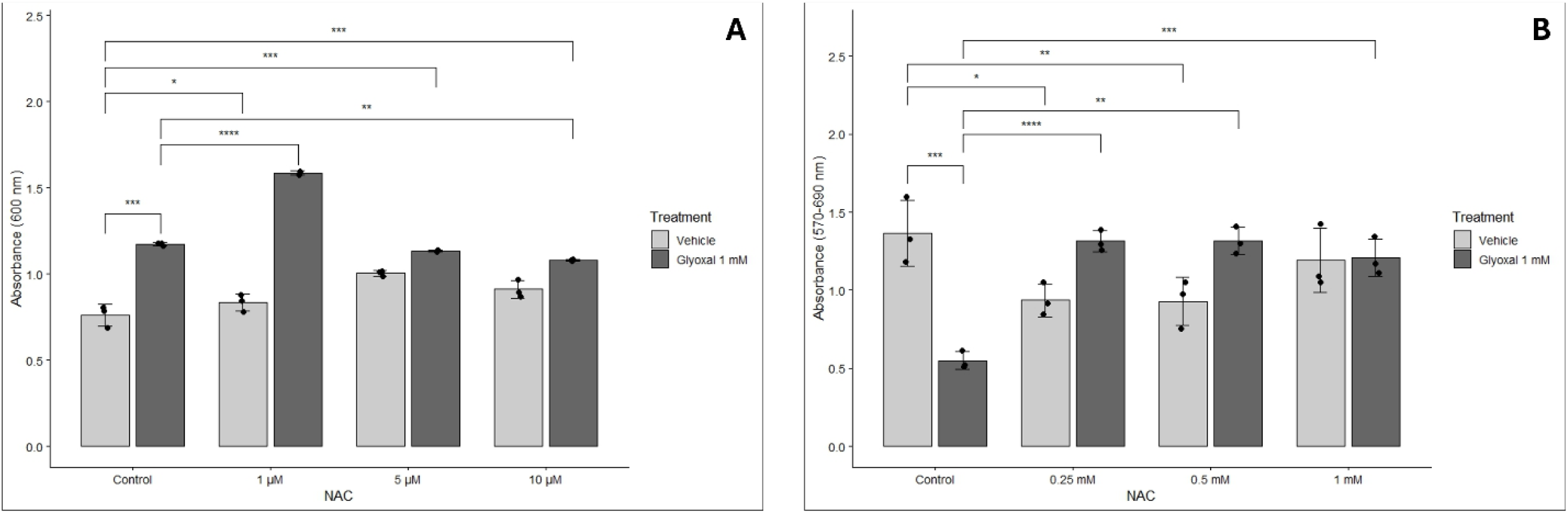
NAC attenuates glyoxal-induced metabolic and redox damage in skeletal muscle. C2C12s were treated with glyoxal at the indicated concentrations in the absence or presence of N-acetylcysteine (NAC) (0.25mM, 0.5mM,1mM). Following treatment, cellular redox status was measured using Alamar Blue reagent **(A)**, and cellular metabolic activity was assessed by MTT reduction **(B)**, as described in Materials and Methods. Data represent means ± SD of triplicate sample measurements. Compared to untreated cells and glyoxal-treated cells, * = significance at p < 0.05, ** = significance at p < 0.01, *** = significance at p < 0.001.

Furthermore, cytoskeletal protection was also observed at NAC 1 mM. Figure 14 shows controls as elongated and proliferating, NAC with similar morphology, methylglyoxal 1mM resulting in rounded and less cells, and the co-treatment showing the return of healthy and dividing cells.

**Figure 14.**
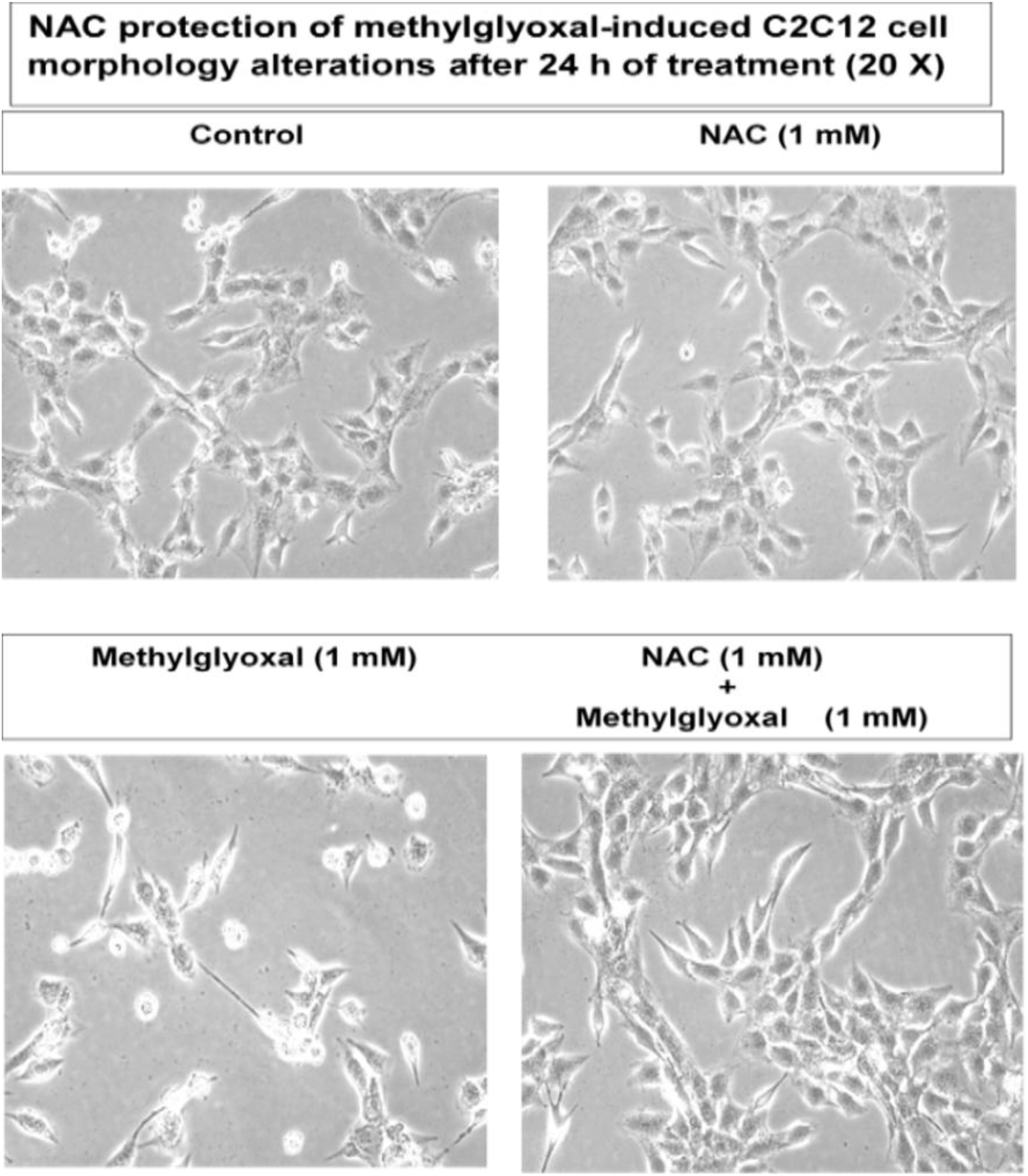
NAC protects skeletal muscle from cytoskeletal degradation. C2C12s were treated with methylglyoxal (1mM) in the absence or presence of NAC 1mMFollowing treatment, images were captured at 20X magnification.

#### Ibuprofen Ameliorates Metabolic Injury in Skeletal Muscle

In response to clear PLA_2_ activation, cells were treated with the COX inhibitor Ibuprofen at varying doses (50, 100, 250 µM) in the absence or presence of glyoxal 1 mM and the metabolic functionality was measured via MTT reduction. Results in figure 15 demonstrated a drastic (98.4%) reduction in metabolic activity in glyoxal 1 mM alone consistent with previous results, and a 51.2% decrease in metabolic activity at the highest dose of Ibuprofen suggesting toxicity. No significant protection results were observed at p < 0.05.

**Figure 15.**
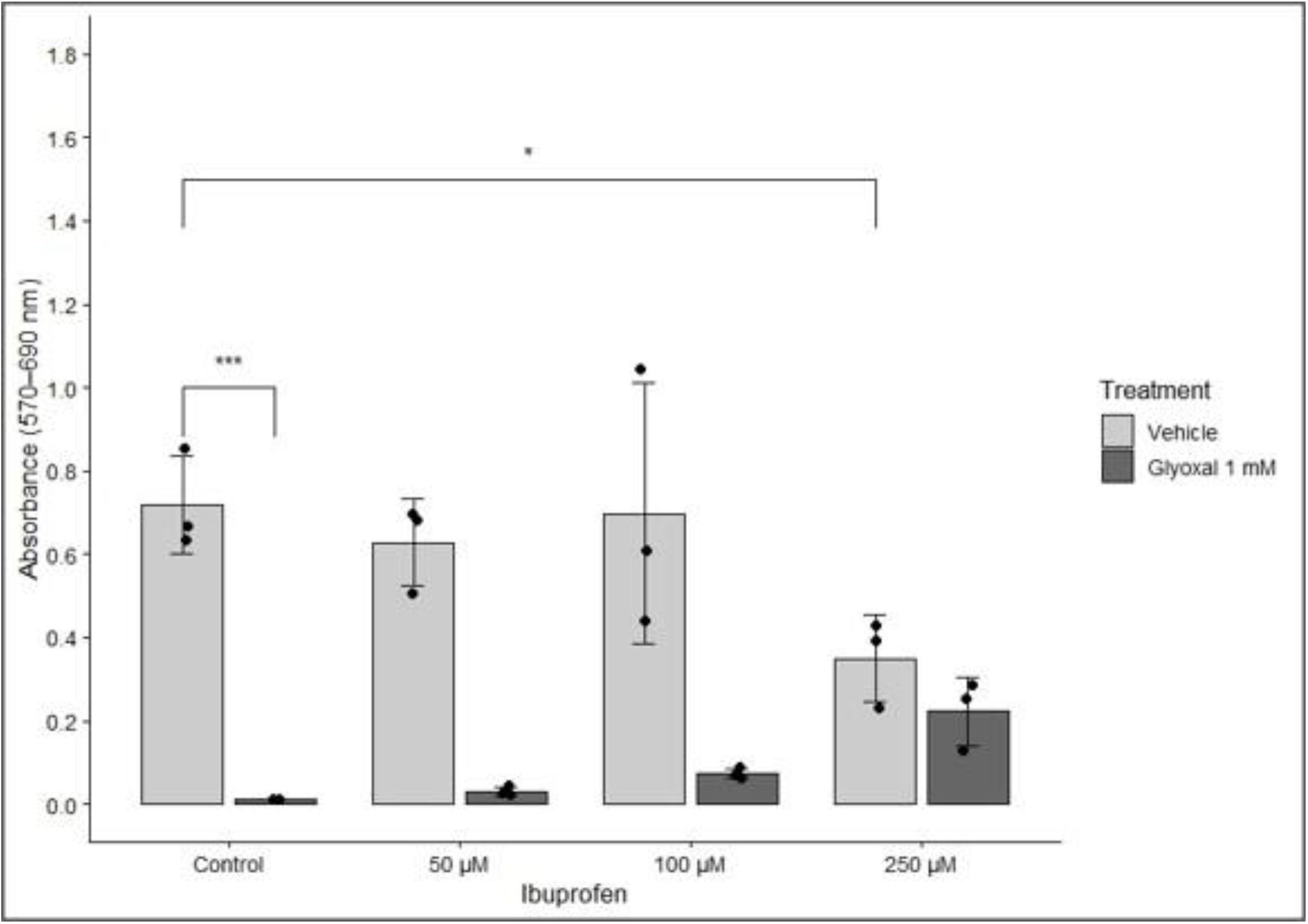
Ibuprofen attenuates glyoxal-induced metabolic dysfunction. C2C12s were treated with glyoxal at 1mM in the absence or presence of ibuprofen (50µM, 100µM, 250µM). Following treatment, cellular metabolic activity was assessed by MTT reduction, as described previously. Data represent means ± SD of triplicate sample measurements. Compared to untreated cells and glyoxal-treated cells, * = significance at p < 0.05, ** = significance at p < 0.01, *** = significance at p < 0.001.

### Inhibition of PLA_2_ Confirms a Possible Mechanism of Action

To further confirm the action of PLA_2_ in diabetic skeletal muscle pathology, cells were treated with varying doses of AACOCF3 (5, 10, 25 µM), a direct PLA_2_ inhibitor, in the absence or presence of glyoxal 1 mM and metabolic activity was measured (Figure 16). Results demonstrated significant protection by the inhibitor of up to 6.4-fold increase compared to glyoxal alone.

**Figure 16.**
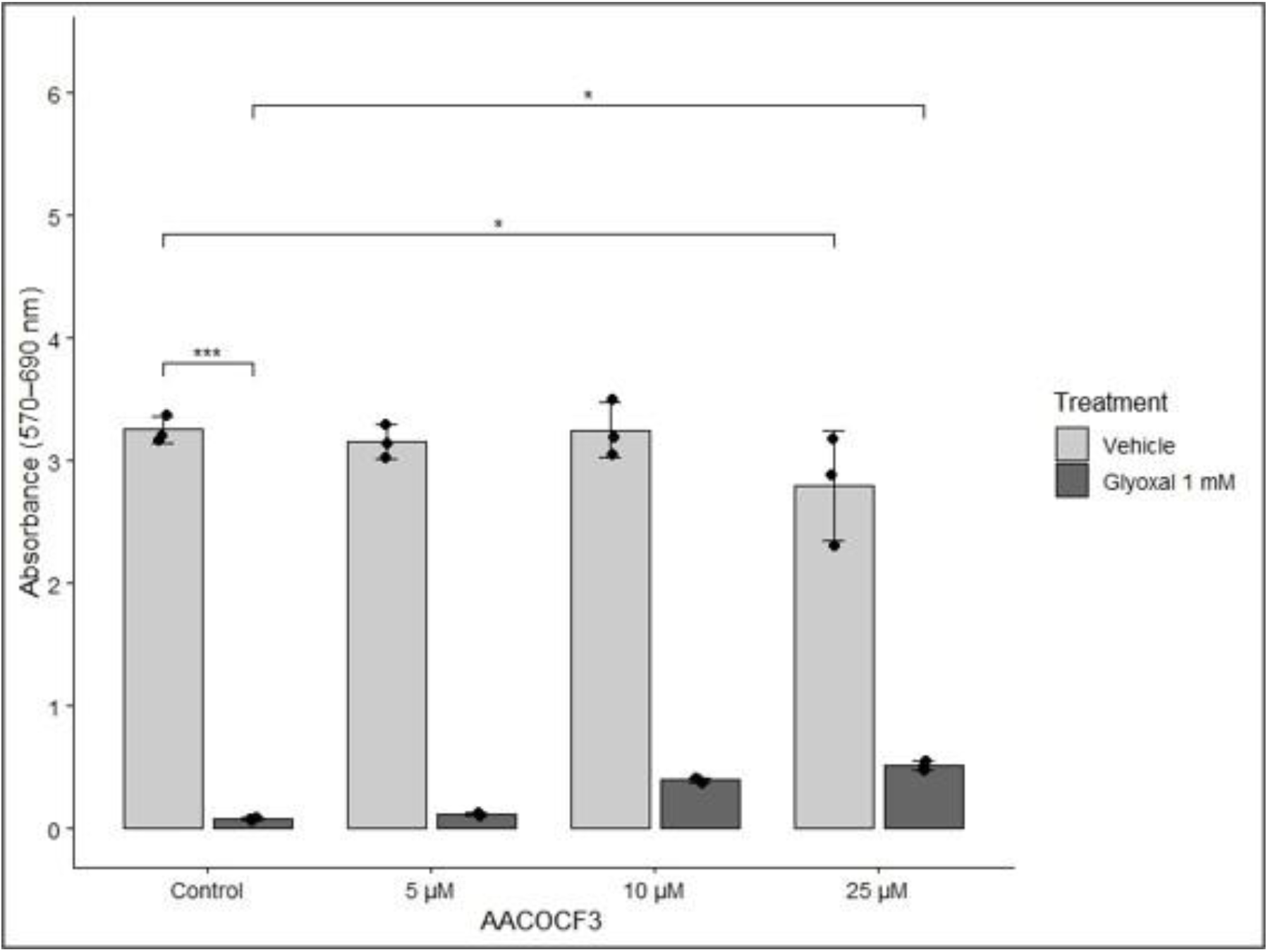
Direct PLA_2_ inhibitor AACOCF3 protects from glyoxal-induced metabolic dysfunction. C2C12s were treated with glyoxal at 1mM in the absence or presence of the phospholipase A2 inhibitor AACOCF3 (5µM, 10µM, 25µM). Following treatment, cellular metabolic activity was assessed by MTT reduction. Data represent means ± SD of triplicate sample measurements. Compared to untreated cells and glyoxal-treated cells, * = significance at p < 0.05, ** = significance at p < 0.01, *** = significance at p < 0.001.

### Metformin Significantly Attenuated Glycolaldehyde-induced Damages

As a current diabetic drug on the market, cells were treated with varying doses of metformin (1,5, 10 mM) to assess attenuation of metabolic loss, cytoskeletal alterations, and PLA_2_ activation. Metformin demonstrated a significant increase in metabolic activity compared to glyoxal challenge of up to a 5.8-fold increase (figure 17A). Furthermore, cytoskeletal alterations were also significantly attenuated by metformin in both bright field morphological visualizations (figure 17B) and in the maintenance of actin fibers in the presence of glyoxal challenge (figure 18A-B). The most significant protection was observed when cells were both pre-treated and co-treated with metformin 5mM in the presence of glyoxal 1 mM (figure 18B)

## Discussion

The present study shows that both MGO and GO cause decreased cellular proliferation, lowered metabolic functioning, decreased redox status, and cytoskeletal alterations in skeletal muscle myoblasts, C2C12s. This is the first description of a mechanism of action within this particular model system suggesting the involvement of various key players in the pathology. Within the model system, treatment with either agonist resulted in increased levels of ROS production contributing to oxidative stress within the cells. Furthermore, a drastic decrease in ATP levels were observed suggesting possible mitochondrial dysfunction further supporting findings of decreased cellular metabolism. Due to suspected mitochondrial injury, a lipid within the inner mitochondrial matrix, cardiolipin (CL), was measured and a dose-dependent decrease was observed. The findings suggest that the reactive glycolaldehydes, MGO and GO, cause the hydrolysis of CL from the mitochondrial membrane producing injury and subsequent mitochondrial dysfunction and metabolic loss.

The mechanism by which this hydrolysis occurs can be attributed to the activation of cPLA_2_. Study observations showed rapid, within 30 minutes, activation of cPLA_2_ when treated with glyoxal at 1 mM. Findings demonstrate an inflammatory characteristic of MGO and GO as well as evidence that CL hydrolysis occurs through MGO/GO-mediated cPLA_2_ activation. Liu et al. found that cardiolipin, due to its high susceptibility to peroxidative damage, could be hydrolyzed by both ROS and cPLA_2_ action in Sprague-Dawley rats. ^22^ Therefore, due to the presence of increased cPLA_2_ activity and ROS production in C2C12s, a plausible mechanism for metabolic injury is supported.

Additional studies have also identified a relationship between cPLA_2_ and the androgen receptor (AR) ^23^. The androgen receptor controls various genes, one of which modulates skeletal muscle mass. In this study, MGO was found to down regulate AR thereby increasing ROS and cPLA_2_ activity. This offers an additional pathway by which ROS, inflammation, and CL loss can occur. Furthermore, due to the inflammatory nature of the PLA_2_ pathway, various inflammation markers such as prostaglandins were measured. Results in figures 11A-H showed a dose dependent increase in PGE1, PGE2, 15-keto PGE2, PGA2, PGD2, Δ12-PGJ2, PGF2α, and 16-keto PGF1α through the activation of cyclooxygenase (COX).

In addition to ROS production, the cytoskeletal degradation observed in figure 5E and figure 18 can be exacerbated by the loss of ATP as shown in figure 8 as the cell loses the energy required to assemble and maintain the cytoskeletal structure. Following the establishment of MGO/GO-induced skeletal muscle damage, points of therapeutic intervention were identified and tested through the use of the antioxidant N-acetyl cysteine (NAC); the COX inhibitors Ibuprofen; and the PLA_2_ inhibitor AACOCF3. NAC demonstrated significant protection of mitochondrial function and redox status at 1 µM but soon dropped at higher doses 5 µM and 10 µM suggesting toxicity from NAC alone at higher doses. This observation demonstrates the utility of NAC as an antioxidant and ROS scavenger but within a strict dosing range. The COX inhibitor Ibuprofen showed a promising but not significant trend in protection for the first two doses of 0.25 mM and 0.5 mM. Future use of inhibitors of this nature would require further toxicity testing to determine if an effective dose is realistic. Inhibition of PLA_2_ by AACOCF3 showed limited toxicity except at the highest dose 250 µM but also demonstrated significant protection at the same dose. This observation aids in supporting our proposed mechanism of glycolaldehyde-induced PLA_2_ activation but holds limited clinical significance due to the housekeeping nature of the enzyme ^24^.

Due to its effectiveness as a drug in cancer, neurodegenerative diseases, and its current use as a diabetic drug, skeletal muscle cells were treated with metformin in the presence of MGO or GO ^25,26^. Although metformin is currently FDA approved for aiding in glucose uptake for diabetic patients, it was observed, as shown in figures 17-18, that it also attenuates loss of metabolism, and restores cell membrane integrity. Effectiveness was best observed at the highest dose, 10 mM, and no significant toxicity was observed, making this drug an ideal candidate to address diabetic-induced muscle weakness or sarcopenia. Based on previous studies by Kinsky et al., metformin can act as a direct scavenger of methylglyoxal, and its compound was isolated from the urine of type-2 diabetic mellitus patients. ^27^ The specific metformin-glyoxal compound was not isolated due to its lower concentration found in the body compared to methylglyoxal, but based on structural similarities, as shown in figure 19, metformin is hypothesized to scavenge both compounds directly. Further study is needed to determine the breadth of metformin action, as it is a vast subject area, but it offers promising results for diabetic patients already prescribed it.

**Figure 17.**
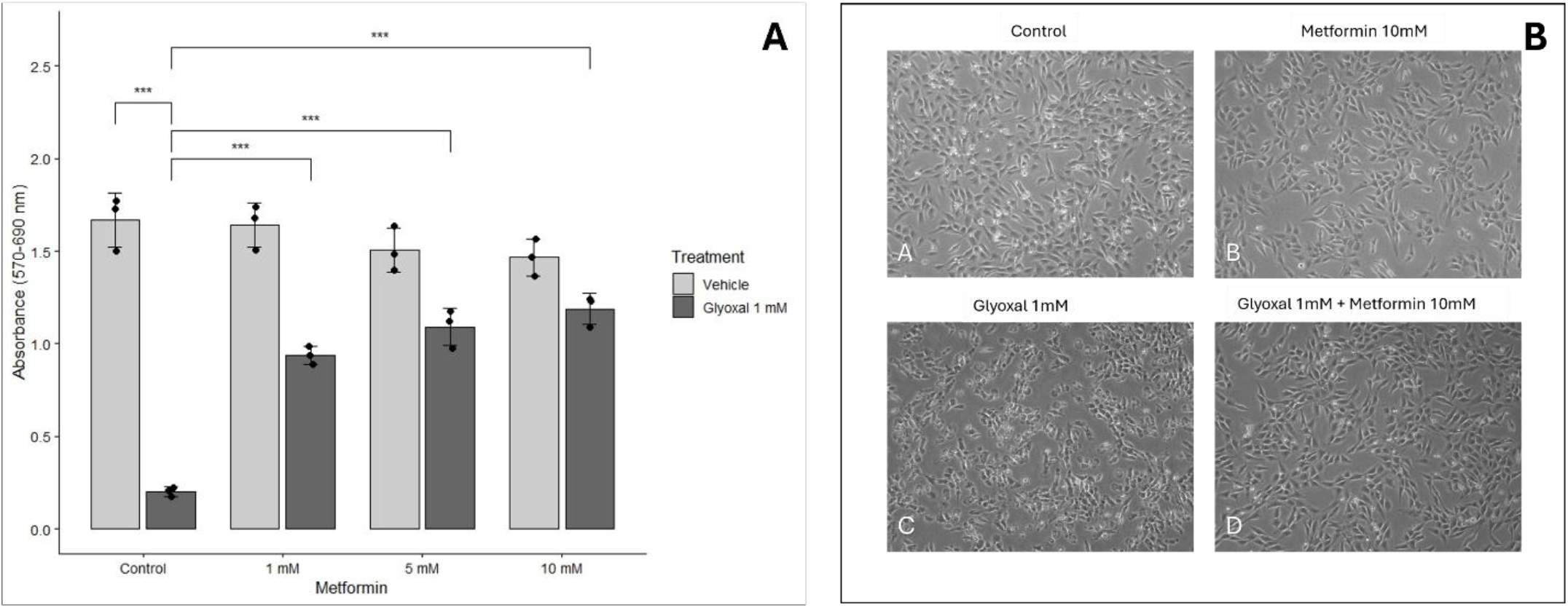
Metformin significantly attenuates glyoxal-induced cytoskeletal alterations and metabolic dysfunction. C2C12s were treated with glyoxal 1mM in the absence or presence of metformin (1mM, 5mM, 10mM). Following treatment, cellular metabolic activity was assessed by MTT reduction **(A)**, and cytoskeletal alterations were assessed by brightfield imaging **(B).** Data represent means ± SD of triplicate sample measurements for quantitative data. Compared to untreated cells and glyoxal-treated cells, * = significance at p < 0.05, ** = significance at p < 0.01, *** = significance at p < 0.001.

**Figure 18.**
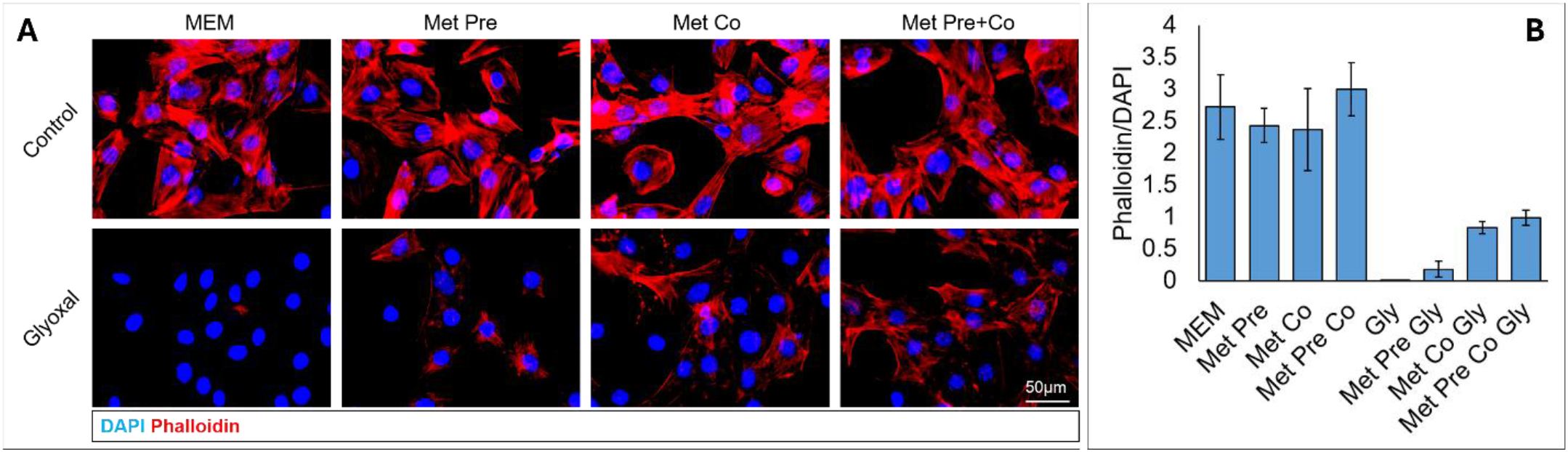
Metformin protects from glyoxal-induced cytoskeletal degradation. C2C12s (70–80% confluency in 96-well culture plates) were treated with glyoxal (1 mM) in the absence or presence of metformin (5 mM). Following treatment, cells were fixed, permeabilized, and stained with phalloidin to visualize the actin cytoskeleton and counterstained with DAPI to visualize nuclei. Fluorescent images were captured at 20X magnification **(A)**, and fluorescence values were quantified using Fiji/ImageJ **(B)**, as described in Materials and Methods. Data represent means ± SD. Compared to untreated cells and glyoxal-treated cells, * = significance at p < 0.05, ** = significance at p < 0.01, *** = significance at p < 0.001.

**Figure 19.**
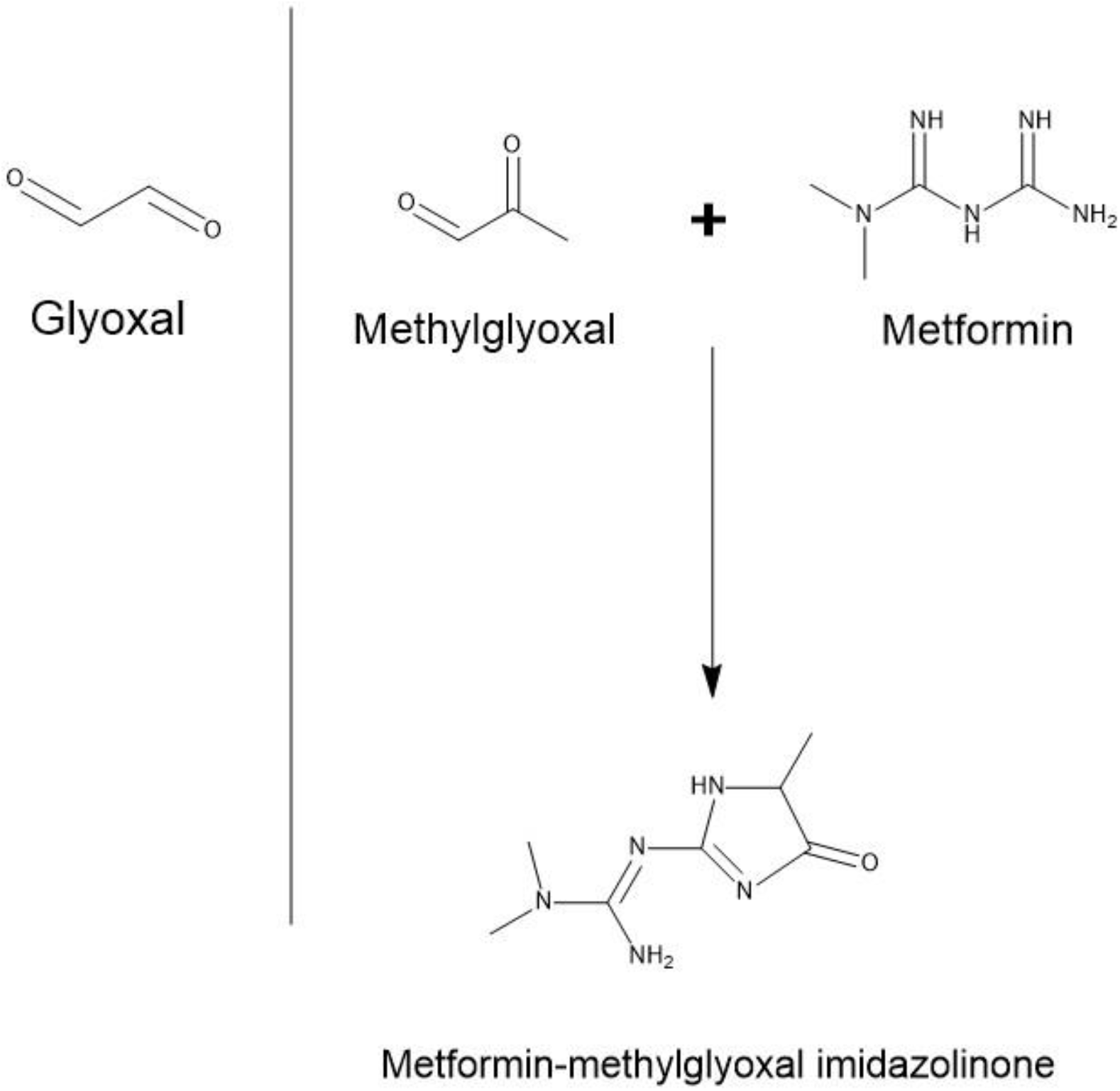
Metformin directly scavenges methylglyoxal. Due to its reactivity metformin can directly scavenge methylglyoxal yielding the compound Metformin-methylglyoxal imidazoline which can be excreted through the urine.

In conclusion, this study showed for the first time that methylglyoxal and glyoxal degrade the cytoskeleton and change the morphology of C2C12 skeletal muscle cells through the formation of AGEs, increased ROS production, and decreased ATP levels; mitochondrial function is lost due to cPLA_2_ activation, increased ROS production, and cardiolipin loss; and inflammation is increased via the PLA_2_ activation of COX and LOX (figure 20). The observed damage can then be attenuated by metformin, COX inhibitors, and NAC at nontoxic doses opening up possibilities of therapeutic intervention for those afflicted with muscle complications.

**Figure 20.**
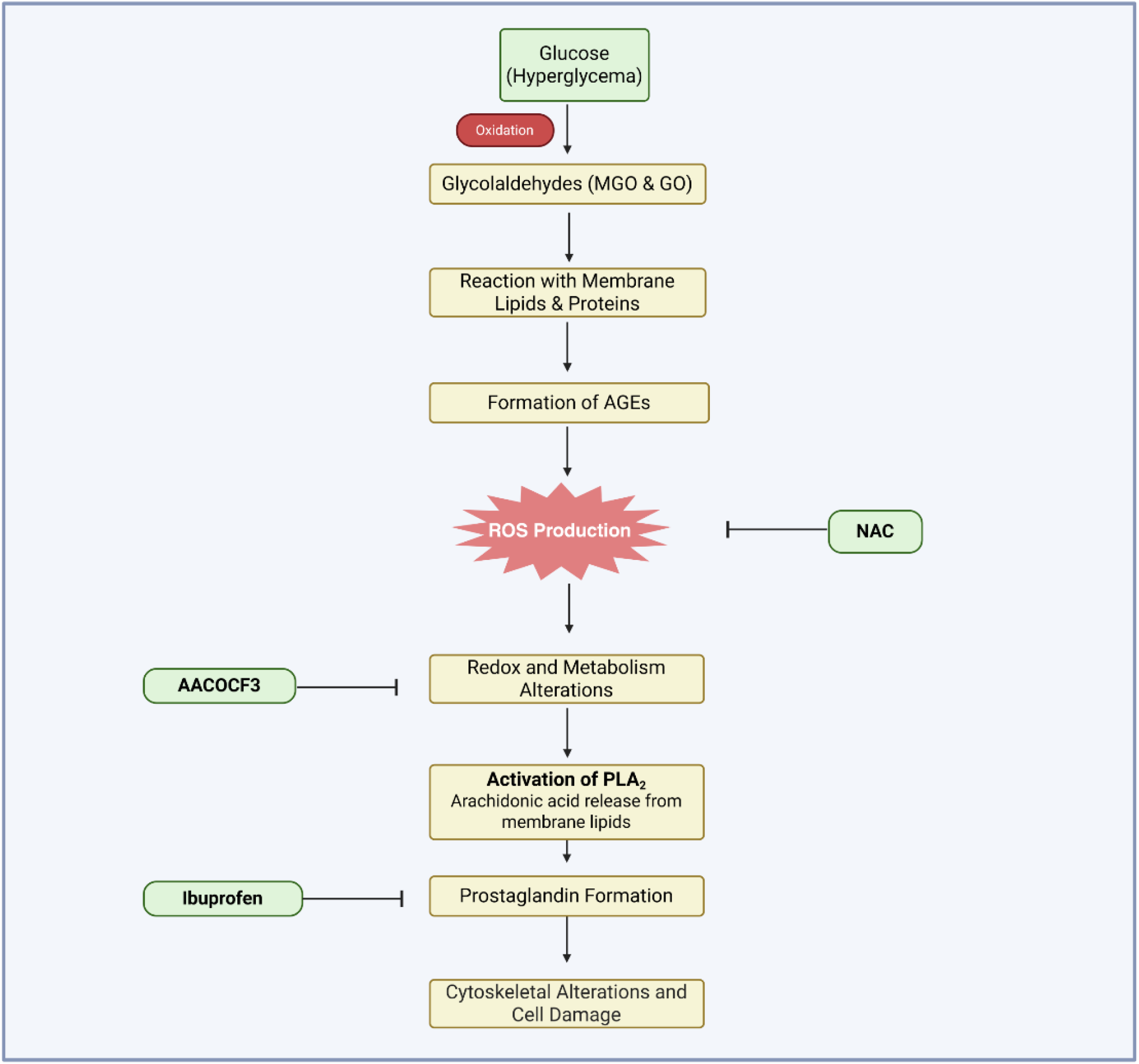
Proposed mechanism for glycolaldehyde-induced skeletal muscle damage. Hyperglycemia promotes the formation of MGO and GO through oxidation steps. The reactive glycolaldehydes react with membrane lipids and proteins to form advanced glycation end products. Elevated ROS levels result and contribute to redox and metabolism alterations, followed by the activation of PLA_2_, prostaglandin formation, and ultimately cytoskeletal alterations and cellular damage. N-acetylcysteine (NAC) attenuates ROS production, whereas AACOCF3 and Ibuprofen inhibit the activation of PLA_2_ and prostaglandin formation respectively.

## Abbreviations

AGEs: Advanced Glycation End Products
AR: Androgen Receptor
ATCC: American Type Culture Collection
BSA: Bovine serum albumin
CEL: Nε-(1-carboxyethyl)lysine
CL: Cardiolipin
CML: Nε-(carboxymethyl)lysine
COX: Cyclooxygenase
cPLA_2_: cytosolic phospholipase A_2_
DCF: 2′,7′-dichlorofluorescein
DCFDA: 2,7- dichlorodihydroxyfluorescein diacetate dicarboxy methyl ester
DMEM: Dulbecco’s Modified Eagle Medium
FBS: Fetal bovine serum GO Glyoxal
LDH: Lactic acid dehydrogenase
LOX: Lipoxygenase
MEM: Minimum essential medium
MGO: Methylglyoxal
MTT: 3-(4,5-dimethylthiazol-2-yl)-2,5-diphenyltetrazolium bromide
NAC: N-acetylcysteine
NAO: 10-N-nonyl acridine orange
PBS: Phosphate buffered saline
PBST: Phosphate buffered saline tween
ROS: Reactive oxygen species

